# Interneuron theta phase locking controls seizure susceptibility

**DOI:** 10.1101/2025.09.10.675457

**Authors:** Zoé Christenson Wick, Paul A Philipsberg, Cassidy Kohler, Sophia I Lamsifer, Elizabeth Katanov, Christopher D Adam, Kathryn E Gordon, Yu Feng, Lauren M Vetere, Genevra C Donnelly, Corin Humphrey, Denise J Cai, Tristan Shuman

**Affiliations:** Nash Family Department of Neuroscience, Icahn School of Medicine at Mount Sinai; New York, 10029, United States

## Abstract

The timing of neuronal activity is highly precise and often organized by brain-wide oscillations. Many neurons modulate their firing rates at specific phases of theta (known as theta phase locking), creating discrete windows for information processing. Disrupted theta phase locking has been found across several neurological and psychiatric disorders (e.g., epilepsy), but gaps in technology have prevented its causal influence from being tested. Here, we developed PhaSER, a closed-loop optogenetic system designed to control the phase locking of specific interneurons, and demonstrate a causal role for inhibitory phase locking in seizure susceptibility. We first found that parvalbumin (PV+) and somatostatin (SOM+) expressing interneurons in the dentate gyrus (DG) show distinct theta phase locking profiles and are differentially impacted in a mouse model of chronic temporal lobe epilepsy. In healthy mice, PV+ interneurons have extremely consistent phase-locked firing near the trough of CA1 theta, aligned with excitatory inputs to DG. However, in epileptic mice, PV+ interneuron activity is dispersed across the theta cycle, suggesting that altered inhibitory phase locking could be a causal mediator of seizure susceptibility in epilepsy. To test this hypothesis, we applied PhaSER to directly control the phase locking of DG interneurons during an acute test of seizure susceptibility. In epileptic mice, re-aligning DG PV+ interneuron theta phase locking reduced seizure susceptibility, while in healthy mice, disrupting normal phase locking of PV+ interneurons increased seizure susceptibility. Together, this provides the first causal evidence that inhibitory theta phase locking can directly control network function by shifting seizure susceptibility in the healthy and epileptic brain.

## INTRODUCTION

The brain relies on precise temporal organization of neuronal spiking to control behavior. Across the brain, network-wide coordination of neural activity is often rhythmic, forming oscillations (e.g., theta) which organize the activity of neurons within and across brain regions^1,2^. This rhythmic synchronization leads many neurons to be active at specific oscillatory phases (i.e., “theta phase-locked”) and creates temporal windows where co-activity can bind neural ensembles and drive distinct cognitive processes^3–8^. Inhibitory neurons are particularly highly phase-locked to theta oscillations, with distinct interneuron subtypes active at different phases^9–14^. This is thought to control the routing of inputs and outputs and maintain excitatory-inhibitory homeostasis. As such, disorganized inhibitory spike timing may impair cognitive computations and disrupt the balance of excitation and inhibition. Yet the only evidence for the importance of phase locking is correlational – extrapolated from situations in which it is disrupted. For instance, phase-locked spiking is altered in models of substance use, schizophrenia, autism, Alzheimer’s disease, and epilepsy^15–23^, suggesting that it may be a common driver of disrupted processing across neurological diseases. Together, this strongly suggests that disruptions to phase locking may be sufficient to induce network dysfunction and alter behavior, but (due to technical limitations) this has never been causally tested.

Under healthy conditions, the dentate gyrus (DG) acts as a “gate” to the hippocampus, with powerful local inhibition selecting which excitatory signals get relayed to downstream regions^24–26^. Waves of rhythmic excitatory inputs enter the DG, carrying spatial information and driving local theta^27,28^. Meanwhile, phase-locked interneurons provide rhythmic inhibition to match these excitatory inputs^16,17^. This puts DG interneurons in a critical position where they must prevent excess excitation from overrunning the network and generating hippocampal seizures^29,30^ – a hallmark of temporal lobe epilepsy (TLE). TLE is the most common form of epilepsy in adults and is often resistant to anti-epileptic drugs. Since most anti-epileptic drugs boost inhibition indiscriminately, this pharmaco-resistance suggests that the mechanisms of seizures are more complex than a bulk imbalance of excitation and inhibition. In fact, using a mouse model of drug-resistant TLE (pilocarpine-induced status epilepticus), previous work has shown a breakdown in inhibitory theta phase locking in the DG of epileptic mice^16^, while major excitatory inputs to the DG maintain their rhythmicity^17^. This suggests that there is a mismatch between the precise timing of excitatory inputs to the DG and local interneuron spiking. We predicted that this breakdown in inhibitory theta phase locking that occurs in the epileptic DG may be a direct causal driver of seizure susceptibility in epilepsy.

To directly test the causal role of inhibitory theta phase locking, we developed and validated an open-source tool, PhaSER (Phase-locked Stimulation to Endogenous Rhythms), that can directly manipulate the phase locking of neurons during behavior. PhaSER performs low-latency theta phase detection and phase-locked optogenetic manipulations, while permitting simultaneous monitoring of spike timing. We used PhaSER to apply phase-specific optogenetic manipulations and directly test the hypothesis that inhibitory phase locking in the DG is a causal mediator of seizure susceptibility in both healthy and epileptic mice.

## RESULTS

### Dentate gyrus PV+ interneurons provide highly precise inhibition at the trough of theta

Very little is known about the *in vivo* firing patterns of distinct subtypes of interneurons in the DG of healthy or epileptic mice. Two prominent and non-overlapping inhibitory cell types in the DG are the parvalbumin (PV+) and somatostatin (SOM+) expressing cells, which have distinct spatial connectivity profiles and spike timing properties^31–35^. While the *in vivo* firing patterns, including theta phase locking, of PV+ and SOM+ cells in the DG are not thoroughly characterized (in either healthy or epileptic animals), studies have shown their intrinsic properties and connectivity profiles are differentially impacted in epilepsy^16,36–39^. These changes could drastically influence the role these cell populations play in their local network, including in their phase locking profiles. We therefore sought to characterize DG PV+ and SOM+ cell theta phase locking in both healthy and epileptic mice and determine how these cells’ phase-locked spiking may impact seizure susceptibility.

To determine the firing patterns of PV+ and SOM+ cells in the DG of healthy and epileptic mice, we conducted acute silicon probe recordings in awake, head-fixed male and female mice as they navigated a virtual linear environment for water rewards (Fig 1A). To isolate the activity of DG PV+ or SOM+ cells, we used a viral-transgenic optogenetic tagging strategy: a Cre-dependent virus expressing the excitatory opsin channelrhodopsin-2 (ChR2) was injected into the dorsal hippocampus of PV-Cre or SOM-Cre mice (Fig 1A). Single-units were identified as opsin-expressing PV+ or SOM+ neurons if they showed a low-latency (<4ms) increase in spiking in response to blue light pulses (Fig 1A). These units’ spike patterns during a baseline period, before any light stimulations were applied, were then analyzed post-hoc.

**Fig. 1.**
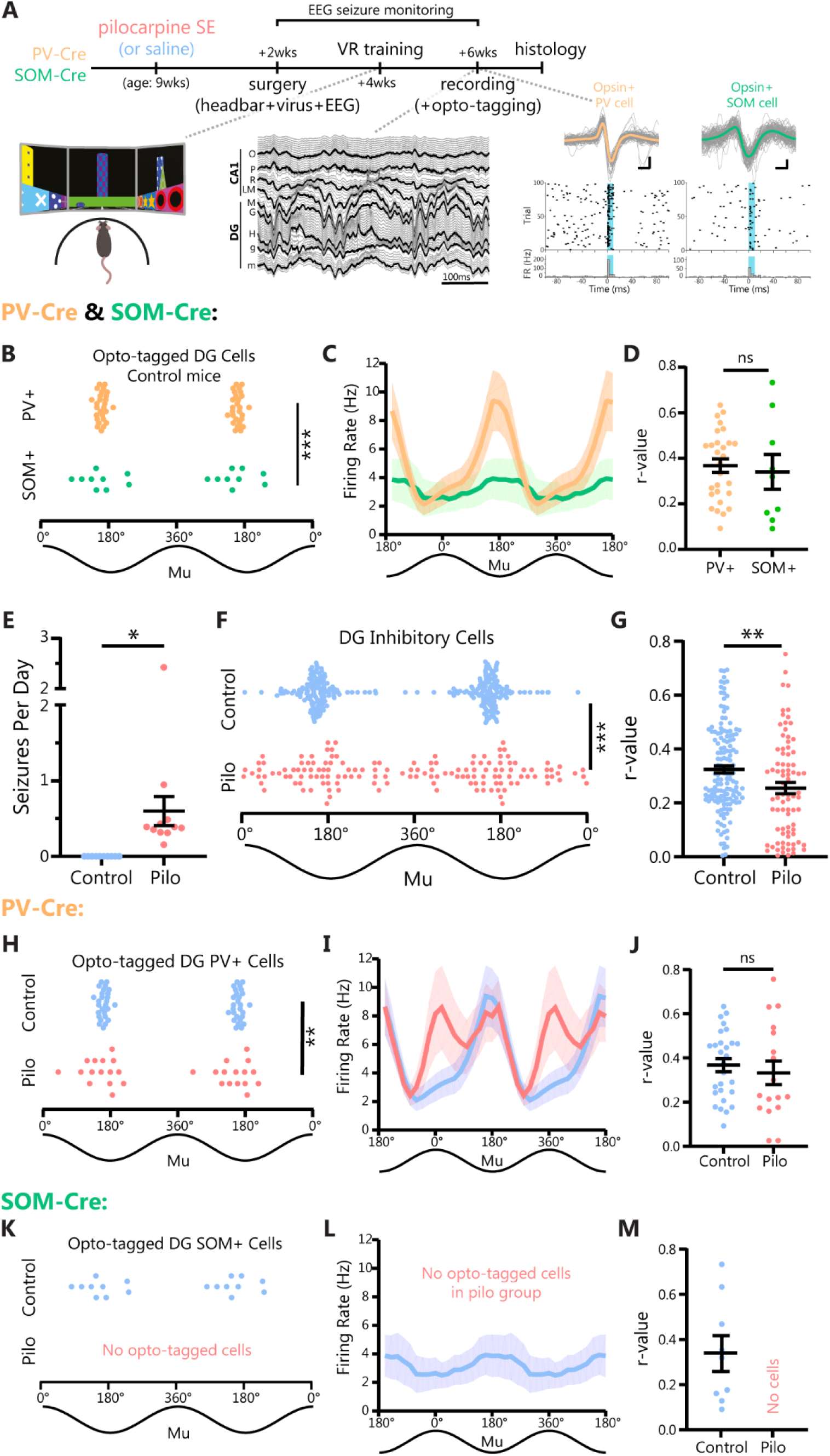
DG PV+ cell firing is tightly locked to the theta trough in healthy mice, and is disrupted in epileptic mice. **A)** PV-Cre and SOM-Cre mice received pilocarpine status epilepticus (SE) or saline (Control), and two weeks later were injected with Cre-dependent ChR2 virus into dorsal hippocampus, and a headbar and wireless EEG transmitter were implanted. Mice were trained to navigate a virtual linear environment and an acute silicon probe recording was performed with channels spanning the dorsal hippocampus. Putative PV+ or SOM+ cells were identified with blue light delivery. Representative waveforms, rasters, and firing rate histograms from a putative opto-tagged PV+ cell (yellow) and SOM+ cell (green). Mean waveforms in yellow and green are shown overlaid against 100 randomly selected individual spike traces from the same cell cluster. Waveform scale: 250µs by 50µV (PV) or 20µV (SOM). **B)** Opto-tagged PV+ cells in the DG of saline-treated control mice showed tightly clustered mean preferred firing phases (mu) near the theta trough. Opto-tagged DG SOM+ cells from saline-treated control SOM-Cre mice also had phase preferences around the theta trough (PV+ vs SOM+ Kuiper test p≤0.05; Watson-Williams test p=0.618), but were significantly more dispersed (PV+ vs SOM+ circular k-test p<0.0001). SOM+: n=9 cells from N=3 mice, PV+: n=28 cells from N=6 mice. **C)** Both DG PV+ and SOM+ cells’ firing rates fluctuated across the theta cycle (20° bins), with firing rates highest in both cell populations around the theta trough, however PV+ cells showed a trend towards greater firing rate modulation based on theta phase. (Two-way repeated measures ANOVA comparing firing based on: cell-type (SOM+ vs PV+) p=0.368, or theta bin p<0.005, or the interaction between cell-type and theta bin p=0.051). **D)** The magnitude of theta phase locking (r-value) was similar in opto-tagged DG PV+ and SOM+ cells in healthy mice (unpaired t-test p=0.685). **E)** Pilocarpine-induced SE produced chronic spontaneous seizures in PV-Cre and SOM-Cre mice (combined), while saline-treated controls did not seize (Welch’s t-test p=0.011, N=10 Control, N=11 Pilo mice). **F)** Inhibitory neurons in the DG of epileptic mice showed altered distribution of mu values (i.e., mean preferred firing phases) relative to controls (Kuiper test p≤0.001; Watson-Williams test p=0.637; circular k-test p<0.0001; Pilo n=72 cells, N=9 mice; Control n=137 cells, N=14 mice). **G)** DG inhibitory neurons in epileptic mice had reduced magnitude of theta phase locking (r-values) compared to controls (unpaired t-test p=0.003; Pilo n=95 cells, N=10 mice; Control n=147 cells; N=14 mice). **H)** Opto-tagged PV+ cells in the DG of epileptic mice had significantly altered distributions of preferred firing phases relative to PV+ cells in controls (Kuiper test p≤0.001, circular k-test p<0.0001) but no shift in the population’s mean firing phase (Watson-Williams test p=0.649; Pilo: n=15 cells, N=3 mice, Control n=27 cells, N=6 mice). Note that the opto-tagged PV+ interneuron data from healthy mice are also shown in panels B-D, and only significantly phase-locked cells are included here. **I)** Opto-tagged PV+ interneurons in the DG showed altered firing rates across the theta cycle (two-way repeated measures ANOVA comparing firing rates based on experimental group (Control vs Pilo p=0.533, theta bin p<0.0001, or the interaction of group and theta bin p=0.021, Pilo n=17 cells, N=3 mice; Control n=28 cells, N=6 mice). **J)** Opto-tagged PV+ cells in the DG showed equivalent strength of theta phase modulation in control and epileptic mice (unpaired t-test, p=0.530; Pilo n=17 cells, N=3 mice; Control n=28 cells, N=6 mice). **K-M)** No opto-tagged SOM+ cells were identified in the DG of epileptic SOM-Cre mice (N=6 Pilo mice), and therefore their theta phase locking profiles could not be characterized. Note that the opto-tagged SOM+ cells from controls are also shown in panels B-D. n=9 cells from N=3 control mice. Note that the theta cycle is double plotted for visualization purposes in panels B, C, F, H, I, K, L. * indicates p<0.05. ** indicates p<0.01. *** indicates p<0.001.

We first examined healthy animals and found that DG PV+ interneurons had extraordinarily tight phase locking to the trough of theta (Fig 1B), with all PV+ interneurons having a mean firing phase (mu) near the trough. This led to a very strong modulation of firing rate relative to the phase of theta, with ∼5x higher firing rates near the trough compared to near the peak of theta (Fig 1C). DG SOM+ neurons also show phase-locked firing near the trough of theta, but the preferred theta phases were much more distributed (Fig 1B), and the population of SOM+ neurons showed notably less overall modulation of spiking across theta phases (Fig 1C). We found no differences in the strength of phase locking for individual neurons (r-values) across PV+ and SOM+ interneurons (Fig 1D), suggesting that the consistency of PV+ cell phase preferences (Fig 1B) is primarily responsible for the increased aggregate phase locking of this population (Fig 1C). Importantly, this precisely trough-aligned spiking matches the spike timing of entorhinal layer 2 neurons, a major excitatory afferent to the DG^17,28^. Thus, in healthy animals, inhibition from PV+ and SOM+ cells is most robust at the trough of theta, when excitatory inputs are strongest. However, we found key differences in the overall phase locking of these populations, with PV+ cell spiking precisely targeted near the trough of theta and SOM+ cell firing more broadly distributed throughout the theta cycle.

### PV+ interneuron theta phase locking is disrupted in epileptic mice

We next explored how the phase locking of PV+ and SOM+ neurons in the DG are impacted in epilepsy. First, we replicated prior work showing that putative inhibitory neurons (Fig S1A,D) in the DG of pilocarpine-treated epileptic mice have substantially altered theta phase locking^16,17^. Specifically, in epileptic mice, inhibitory neurons in the DG had theta phase preferences that were more distributed throughout the theta oscillation (Fig 1E,F, Fig S1B,E), and their firing was less strongly modulated by theta phase (Fig 1G, Fig S1C,F). However, we saw no change in their overall firing rate (Fig S1G,I)^17^.

When looking at opto-tagged DG PV+ and SOM+ cells specifically, we found that DG PV+ cells in epileptic mice had significantly more distributed theta phase preferences than in healthy animals (Fig 1H,I), though there was no change in the strength of their theta phase modulation in individual neurons (r-value; Fig 1J). Thus, the distributed firing activity of the population of PV+ interneurons in epileptic mice (Fig 1H) was driven by the population having much more variable preferred phases of firing, rather than each neuron’s spiking being more distributed. We saw no overall change in PV+ cell firing rates (Fig S1H). Meanwhile, we were unable to identify any DG SOM+ cells in epileptic animals using optogenetic-tagging strategies. This is likely due to the substantial SOM+ cell loss that occurs in temporal lobe epilepsy^16,40–43^ and the low throughput of opto-tagging. Altogether, while the majority of DG SOM+ cells are lost in TLE, the remaining interneurons, including PV+ cells, show more distributed theta phase locking.

Importantly, we confirmed that epileptogenesis was equally effective at producing spontaneous seizures in both PV-Cre and SOM-Cre mouse strains (Fig S1K), and saw virtually identical deficits in overall DG inhibitory theta phase locking in both strains as well (Fig S1B,C,E,F). However, there was a notable sex effect with the phase locking deficit primarily occurring in males (Fig S2A,B). Meanwhile, DG inhibitory neurons in epileptic females showed theta phase locking profiles that were more similar to healthy females (Fig S2A,B,D). Interestingly, this sex-specific change in theta phase locking occurred despite males and females having similar seizure frequencies (Fig S2C). Thus, there may be sex-specific mechanisms driving altered spike timing and seizures in TLE.

The more distributed inhibitory spike timing in epileptic mice (Fig 1F,H) may be a compensatory mechanism to tamp down hyperexcitability, but it also alters the relative distribution of inhibition across the theta cycle. This may disrupt homeostatic plasticity mechanisms^44,45^ and leave the DG vulnerable to hyperexcitability when rhythmic excitatory inputs are strongest (i.e., near the theta trough). Given the theorized importance of inhibitory spike timing in shaping circuit-wide dynamics, and the changes we found in epileptic mice, we hypothesized that this disruption in inhibitory theta phase locking might causally contribute to the emergence of seizures in TLE. To test this hypothesis, we performed direct manipulations of PV+ and SOM+ cell phase locking while measuring seizure susceptibility.

### PhaSER: a tool for real-time theta phase estimation and phase-locked manipulation

To experimentally manipulate phase locking, we developed PhaSER (Phase-locked Stimulation to Endogenous Rhythms), an open-source platform for delivering fast, flexible, and highly accurate closed-loop perturbations at any specified phase of an oscillation during behavior (Fig 2A; Fig S3). To estimate theta phase, PhaSER performs real-time zero-phase filtering for theta on a reference electrode (in our recordings, we referenced the deep pyramidal cell layer of CA1 and phase was estimated every 1ms). However, real-time filtering introduces distortions at the edge of the sampling window that must be trimmed off. An autoregressive forward prediction model ^46^ was therefore trained on each reference signal and then applied during the recording to extrapolate the trimmed/distorted signal. Then, the Hilbert transform was applied for phase estimation, triggering an LED for optogenetic manipulations at any specified phase (Figs S3, S4). We found that PhaSER estimates theta phase with outstanding accuracy, delivering light pulses to specified theta phases precisely and with low-latency (Fig 2B; Fig S4A-C) across a range of physiological theta powers in mice as they traverse a virtual linear track (Fig S4D).

**Fig. 2.**
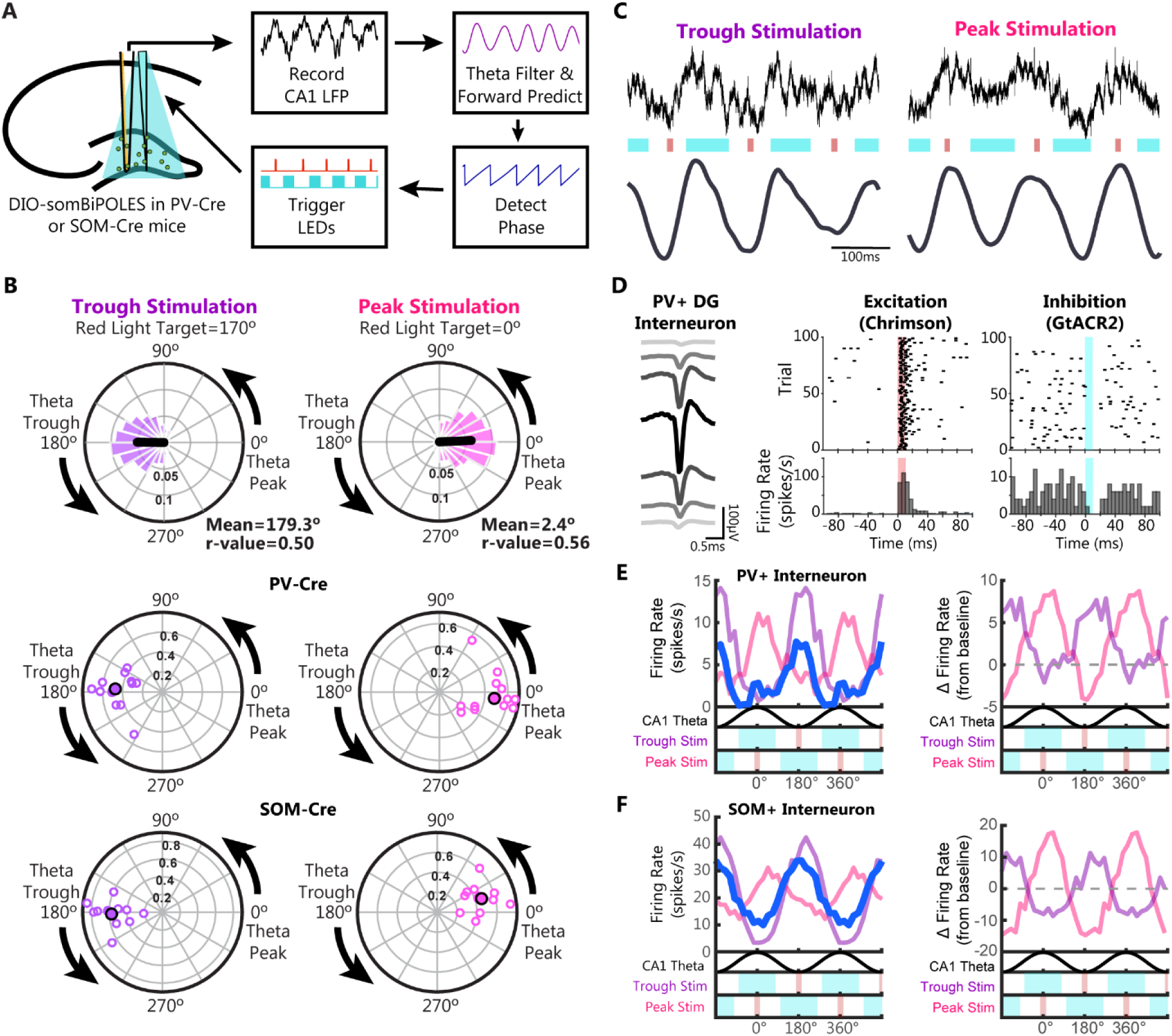
Closed-loop manipulation of inhibitory theta phase locking with PhaSER. **A)** Schematic of PhaSER: a deep pyramidal layer CA1 reference channel is filtered for theta (4-9Hz) before theta phase detection is applied to independently trigger red and blue light delivery at specific theta phases with low latency. See Methods and Fig S3 for details on autoregression used for forward prediction of theta phase. **B)** Top: Circular distribution of red light pulses in a representative animal targeted to the theta trough (left, target 170°) or peak (right, target 0°) showing accurate LED triggers centered around the target phase (mean phase of red light delivery when target is 170° = 179.3°; when target is 0° mean phase = 2.4°) with high precision (trough r-value = 0.50, peak r-value = 0.56). Bottom: polar plot showing the distribution of the mean phase of light delivery across N=13 PV-Cre and N=12 SOM-Cre mice when targeting the trough (in PV-Cre mice: 177.0°, SOM-Cre: 182.8°) and peak (PV: -7.3°, SOM: 16.0°) of CA1 theta. The trigger precision (r-value) is shown on the radial axis (PV trough: 0.47±0.04; PV peak: 0.49±0.04; SOM trough: 0.57±0.05; SOM peak: 0.47±0.04). No difference in precision of trough or peak targeted stimulation in PV-Cre compared to SOM-Cre mice (two-sample t-tests, p>0.05). **C)** Representative traces showing excitatory red light targeted to the trough (left) or peak (right) of theta with inhibitory blue light targeted ± 90° around the opposing phase. Raw signal shown on top, with 5-12Hz theta filtered trace on bottom. **D)** Bidirectional manipulation of spiking in a somBiPOLES expressing (i.e., opto-tagged, or PV+) neuron in the dentate gyrus of a PV-Cre mouse. Left: average spike waveform of optotagged PV+ interneuron across neighboring electrode channels. Right: Red light increases spiking with low latency (<4ms) while blue light suppresses spiking in the same neuron. **E-F)** Mean firing profiles relative to CA1 theta of individual opto-tagged cells from a PV-Cre (E) and SOM-Cre (F) mouse during baseline (blue), trough (purple), and peak (pink) targeted stimulation periods. Peak Stim (red excitatory light targeted to peak, blue inhibitory light delivered around the trough) shifted preferred firing to the peak of theta and reduced firing at the trough, while Trough Stim (red excitatory light targeted to the trough, blue inhibitory light around the peak) enhanced preferred firing at the trough of theta.

### Bidirectional manipulations shift the theta phase preference of PV+ and SOM+ cells

To validate PhaSER’s ability to control theta phase locking, we first tested if applying phase-specific excitatory blue light to ChR2-expressing PV+ or SOM+ cells could alter their theta phase locking profiles. We found that exciting PV+ or SOM+ cells at various theta phases produced additional spikes at the targeted phase but did not reduce firing at the initially preferred phase (Fig S5). We therefore tested whether a bidirectional manipulation, with excitation and inhibition locked to opposing phases of theta, could successfully shift the phase preference of opsin+ neurons. To do this, we expressed a Cre-dependent soma-targeted BiPOLES opsin^47^ in the dorsal hippocampus of PV-Cre and SOM-Cre mice (Fig S6). BiPOLES permits bidirectional manipulation of spiking, with red light inducing spiking via Chrimson, a red-shifted channelrhodopsin variant, and blue light inhibiting spiking via GtACR2, a blue light-activated chloride channel (Fig 2D). To test if this bidirectional manipulation was capable of shifting the phase preference of BiPOLES-expressing PV+ and SOM+ cells, we triggered a 10ms excitatory red light pulse at the target phase (e.g., trough or peak) and triggered a 180° window of inhibitory blue light centered around the opposite phase of theta (e.g., peak or trough, respectively; Fig 2C). This pattern of theta-locked dual light manipulation was indeed capable of shaping the firing profiles of BiPOLES-expressing PV+ and SOM+ cells. In both PV+ (Fig 2E) and SOM+ (Fig 2F) interneurons, peak excitation with trough inhibition (Peak Stim) was able to shift their preferred firing phase to the peak of theta while reducing firing at the trough (Fig 2E-F). Meanwhile trough excitation with peak inhibition (Trough Stim) enhanced their trough preference (Fig 2E-F). Thus, this bidirectional approach was able to fully control the preferred phase of these subpopulations of inhibitory neurons *in vivo* during awake behavior, enabling us to directly test the causal role of theta phase locking in PV+ and SOM+ inhibitory neurons.

### Oscillatory power and theta phase are not altered by phase-specific PV+ or SOM+ cell manipulations

Some inhibitory neurons in the dorsal hippocampus have been shown to contribute to the local generation of theta^48,49^ (but see also ^50,51^) or reset the theta phase upon activation^52^. If our inhibitory phase locking manipulations were to alter the theta oscillation itself, this would not only impact PhaSER’s ability to perform accurate phase estimation but would also impact the interpretation of our experimental results. We therefore tested whether manipulating DG PV+ or SOM+ cell theta phase locking impacted the power or phase of the endogenous oscillatory activity broadly, and in the theta band in particular. To test this, we first compared oscillatory power during running bouts without stimulation (baseline) to running bouts where trough- or peak-locked bidirectional stimulations were applied (Fig 3A,E). Notably, we did not detect a change in local field potential (LFP) power (1-100Hz) in either the DG or CA1 during trough- or peak-locked manipulations of PV+ or SOM+ cell activity (Fig 3A,E; Fig S7). To further confirm that theta was not impacted by these phase-specific manipulations, we compared broadband theta (5-12Hz) in the referenced layer (CA1) during baseline, trough-, or peak-targeted stimulation periods and found no significant change in power when either PV+ cells (Fig 3B) or SOM+ cells (Fig 3F) were manipulated.

**Fig. 3.**
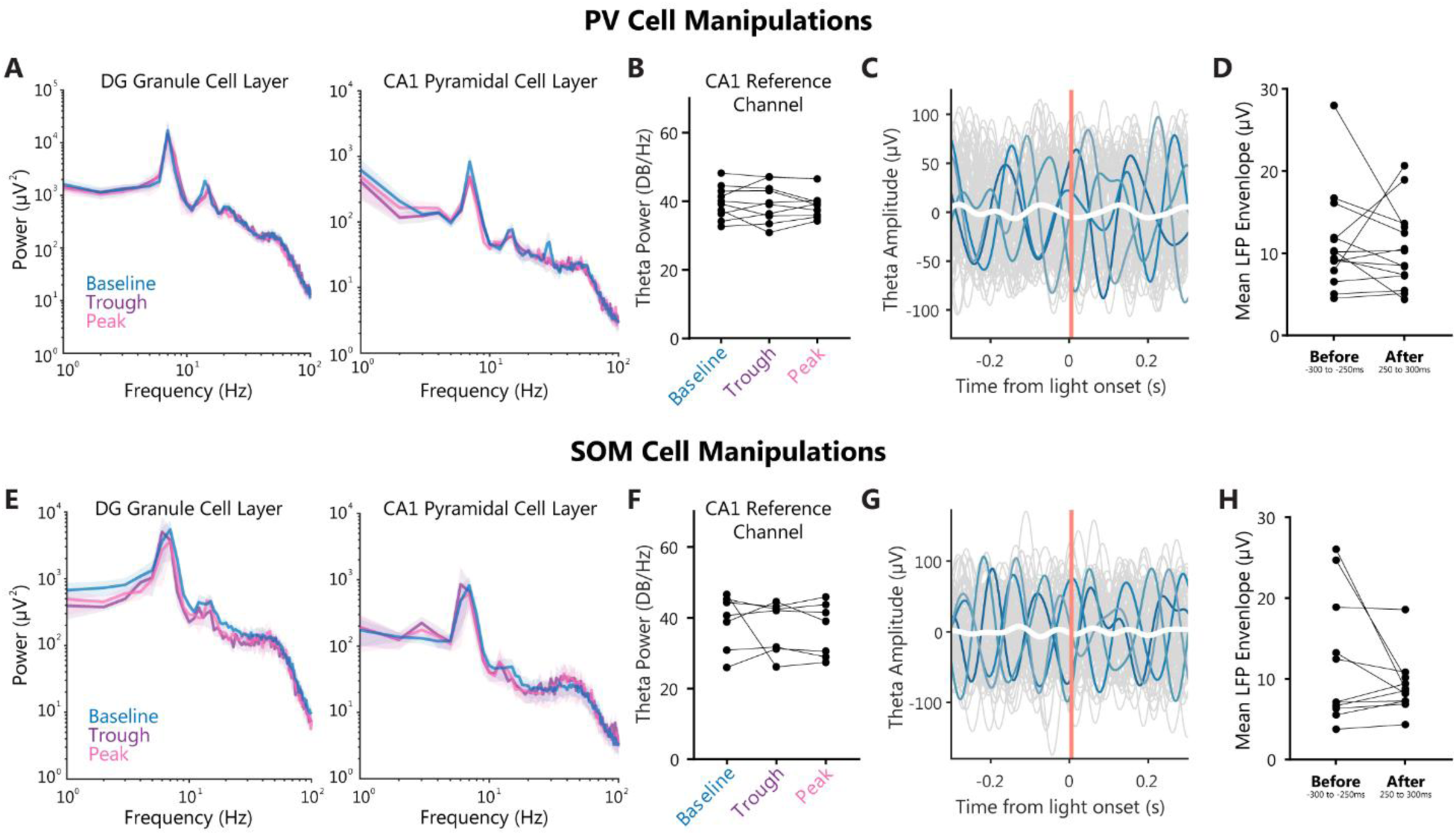
Manipulating dentate PV+ or SOM+ cell theta phase preference does not impact the referenced LFP. **A-H)** No change in oscillatory power or theta phase during DG PV+ cell (A-D) or SOM+ cell (E-H) trough- or peak-targeted manipulations compared to baseline. **A,E)** No changes in power spectral density locally in the DG granule cell layer with PV+ cell theta phase manipulations [A, left; two-way repeated measures ANOVA, no significant effect of manipulation (baseline vs trough-vs peak-stim, p=0.688) and no significant interaction between frequency and manipulation (p=0.279), N=10] or SOM+ cell theta phase manipulations [E, left; two-way repeated measures ANOVA, no significant effect of manipulation (baseline vs trough-vs peak-stim, p=0.609) and no significant interaction between frequency and manipulation (p=0.692), N=5]. No changes in power spectral density in the CA1 pyramidal cell layer with PV+ cell theta phase manipulations [A, right; two-way repeated measures ANOVA, no significant effect of manipulation (baseline vs trough-vs peak-stim, p=0.123) and no significant interaction between frequency and manipulation (p=0.138), N=10] or SOM+ cell manipulations [E, right; two-way repeated measures ANOVA, no significant effect of manipulation (baseline vs trough-vs peak-stim, p=0.123) and no significant interaction between frequency and manipulation (p=0.138), N=10]. **B,F)** No significant change in theta band power (5-12Hz) in the theta reference channel (i.e., CA1 deep pyramidal cell layer) between baseline, trough-, and peak-stimulation conditions when manipulating PV+ cells (B, one-way repeated measures ANOVA, p=0.956, N=10) or SOM+ cells (F, one-way repeated measures ANOVA, p=0.589, N=7). **C,G)** Filtered theta from the CA1 pyramidal cell reference layer from representative animals, aligned to 100 red light pulses (10ms duration) showing no theta phase reset when stimulating PV+ cells (C) or SOM+ cells (G). Randomly selected individual traces are colored in shades of blue, mean of all traces is in white. Note that this signal has been filtered for theta (5-12Hz) for visualization purposes, but analysis in D,H is on unfiltered data. **D,H)** The LFP envelope was calculated from down-sampled, unfiltered data aligned to light pulses, as in C and G, on a 50ms window starting 250ms before or 250ms after light delivery. No significant change in LFP envelope was noted before versus after stimulation of either PV+ cells (paired t-test, p=0.672, N=14) or SOM+ cells (paired t-test, p= 0.189, N=11).

Finally, we tested if stimulating PV+ or SOM+ cells in the DG would reset theta phase. To test for a theta phase reset, we aligned the reference channel’s LFP to excitatory red light pulses (as in opto-tagging protocol). We then compared the envelope of the averaged LFP signal before versus after light delivery. If stimulating PV+ or SOM+ neurons was resetting the theta phase, we would expect to see an increase in the mean LFP envelope lasting more than one theta cycle (i.e., >120ms). However, we did not see a change in the mean envelope when either PV+ cells (Fig 3C,D) or SOM+ cells (Fig 3G,H) were stimulated. Together, these results suggest that manipulating DG PV+ or SOM+ cell theta phase locking does not alter theta power or theta phase in the hippocampus.

### PV+ cell theta phase locking bidirectionally controls seizure susceptibility

Hippocampal theta is generated, in large part, by rhythmic excitatory inputs from the medial entorhinal cortex, including direct inputs from layer 2 stellate cells to the DG^27^. These excitatory inputs are maximally active near the trough of theta in healthy mice and their rhythmic firing is maintained in epileptic mice^17^. In the healthy hippocampus, inhibition is precisely matched to the trough of theta, but in epilepsy DG inhibition is much more dispersed (Fig 1, Figs S1, S2)^16,17^. We predicted that this disruption in the precise timing of inhibition weakens the relative inhibitory strength at the theta trough, making the DG vulnerable to overexcitation and increased seizure susceptibility. We therefore set out to test whether re-aligning DG inhibition to the theta trough in epileptic male mice would reduce seizure susceptibility, and whether shifting DG inhibition away from the theta trough in healthy mice would increase seizure susceptibility (Fig 4A). Due to the relatively low rate of spontaneous seizures (Fig 1E, Figs S1K, S2C), and to allow investigation of seizure susceptibility in healthy mice, we measured seizure susceptibility using an acute intraperitoneal injection of the chemoconvulsant kainic acid (KA) and measured the latency to the first seizure (Fig 4B)^53,54^. We titrated the dosage of KA differently in healthy and epileptic mice so that an acute seizure would be reliably induced within ∼5 minutes in epileptic mice (20mg/kg KA), and within ∼10 minutes in controls (30mg/kg KA). We were then able to determine if re-aligning DG inhibitory spiking to the trough would increase seizure latency in epileptic mice, and if mis-aligning DG inhibitory spiking to the peak might shorten seizure latency in otherwise healthy mice.

**Fig. 4.**
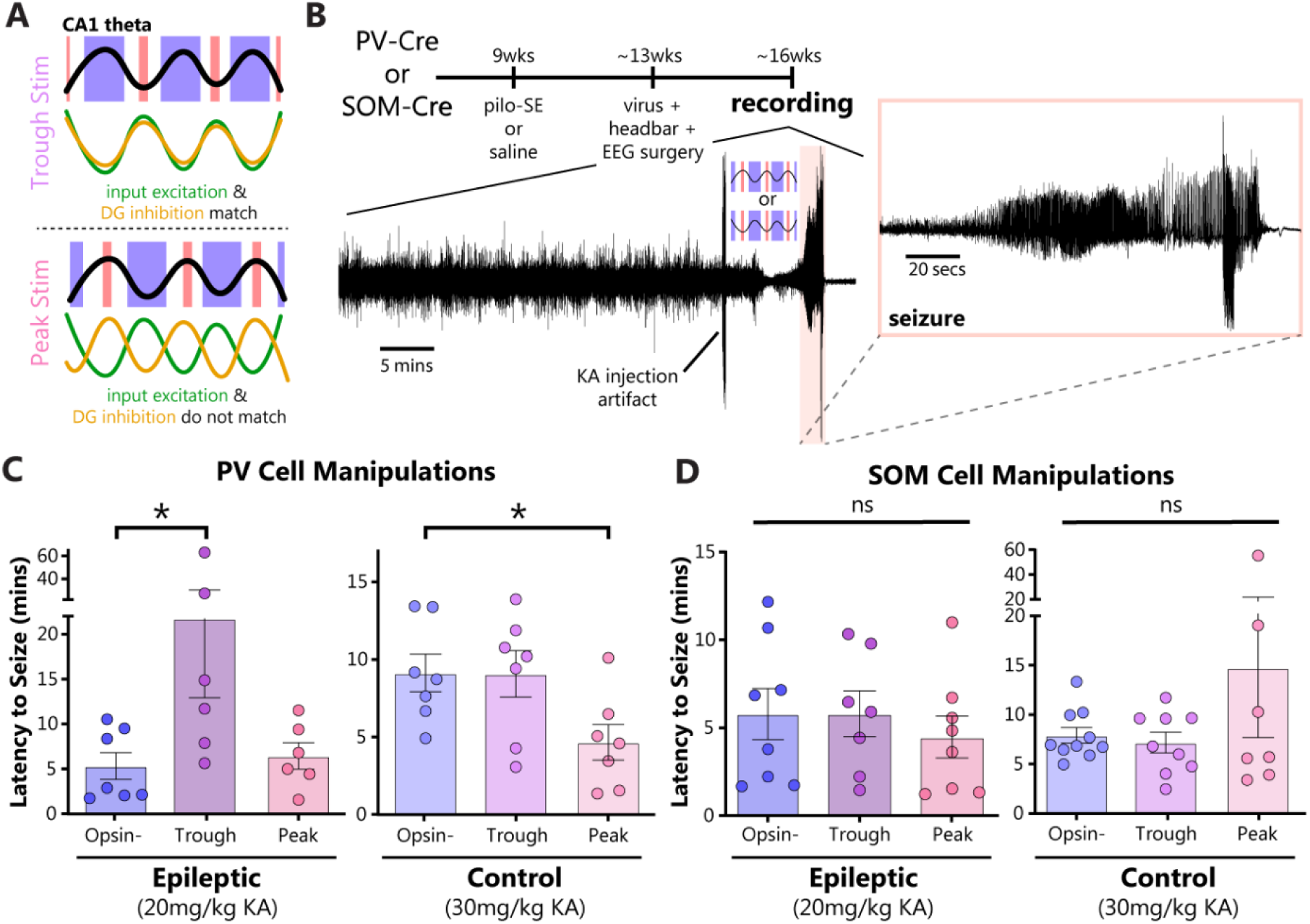
PV+ interneuron theta phase locking impacts seizure susceptibility. **A)** Schematic of hypotheses. Top: in epileptic mice, Trough Stim (trough excitation with peak inhibition) re-aligns DG inhibition to the trough of CA1 theta, when input excitation is strongest. Bottom: in control mice, Peak Stim (peak excitation with trough inhibition) mis-aligns DG inhibition, so inhibition is weakest when input excitation is strongest, creating seizure vulnerability points at the theta trough. **B)** Experimental timeline schematic. PV-Cre and SOM-Cre mice received pilocarpine status epilepticus (SE) or saline (Control), and four weeks later were injected with Cre-dependent somBiPOLES virus into dorsal DG, and a headbar and wireless EEG transmitter were implanted. Three weeks later, an acute silicon probe recording was performed with channels spanning the dorsal hippocampus. Following a baseline period, mice were injected intraperitoneally with kainic acid and either peak-targeted or trough-targeted stimulation was applied until seizure onset. **C)** Latency to seizure in epileptic PV-Cre mice (left, male mice) was significantly increased compared to opsin-when PV+ cells were re-aligned to the trough of CA1 theta (Kruskal-Wallis ANOVA p=0.040; with Dunn’s post hoc tests comparing Trough vs Opsin-, p=0.041; and Peak vs Opsin-, p>0.999). Latency to seizure in PV-Cre control mice (right, male and female mice) was significantly reduced compared to opsin-when PV+ cells were mis-aligned to the peak of CA1 theta (one-way ANOVA p=0.039 with Dunnett’s multiple comparison post hoc tests comparing Trough vs Opsin-, p>0.999; and Peak vs Opsin-, p=0.046). **D)** No significant effects of manipulating SOM+ cell phase locking on latency to seizure in epileptic (left, one-way ANOVA p=0.741) or control (right, one-way ANOVA p=0.273) mice.

In epileptic mice, we found that stimulating DG PV+ cells at the trough of theta (while inhibiting at the peak) significantly increased the latency to KA-induced seizure compared to opsin-negative (opsin-) animals, while stimulating PV+ cells at the peak of theta (and inhibiting at the trough) had no effect on seizure latency (Fig 4C, left). Notably, this effect was only seen when manipulating PV+ cell firing in epileptic male mice, and not in epileptic female mice (Fig S8A), potentially because epileptic females did not have severely disrupted theta phase locking (Fig S2A,B,D). Meanwhile, we found no change in seizure duration with peak or trough stimulation (Fig S8C-D), and neither circadian or multidien rhythms, nor time since epileptogenesis could explain the variability in seizure latency (Fig S8E). Thus, correcting DG PV+ interneuron theta phase locking is sufficient to reduce seizure susceptibility in the epileptic hippocampus.

We next tested if mis-aligning DG PV+ interneuron theta phase locking could exacerbate seizure susceptibility in otherwise healthy animals. Here, we combined sexes as we did not see differences in DG inhibitory theta phase locking between healthy male and female mice (Fig S2A,B; though a breakdown by sex can be found in Fig S8A,B). In healthy mice, we found that mis-aligning DG PV+ cell spiking to the peak of theta significantly reduced the latency to KA-induced seizure compared to opsin-animals, while stimulating PV+ cells at the trough had no effect on seizure latency (Fig 4C, right). Notably, this effect was specific to DG PV+ interneurons as manipulating theta phase locking of DG SOM+ neurons had no effect on seizure latency in healthy or epileptic mice (Fig 4D). While the lack of effect in the epileptic SOM-Cre animals is perhaps expected given the well-documented loss of DG SOM+ cells in TLE (Fig 1K-M)^40–42^ (but see also ^55^), the lack of effect in healthy SOM-Cre animals suggests that SOM+ interneurons have a less powerful role in regulating DG excitability. Altogether, these results demonstrate that PV+ interneurons are a powerful regulator of hippocampal excitatory-inhibitory homeostasis, and PV+ interneuron theta phase locking is a causal mediator of seizure susceptibility.

## DISCUSSION

We found that dentate gyrus PV+ interneuron theta phase locking can bidirectionally control seizure susceptibility in both healthy and epileptic mice. DG inhibitory neurons, particularly PV+ and SOM+ cells, are critical mediators of hippocampal processing, selecting which incoming signals are passed on for further processing^24,32^. In healthy mice, we found that PV+ cells provide highly precise inhibition at the trough of theta, while SOM+ neurons had more dispersed firing across theta phases. This suggests that PV+ interneurons are particularly important for balancing excitatory inputs from MEC that arrive near the trough, while the more dispersed firing of SOM+ interneurons suggests a broader role in dentate function^35,56,57^. In epileptic mice, alterations in DG inhibitory connectivity and function^16,36,37,40,42,55^ are likely to have an avalanche of consequences. Indeed, in epileptic mice, DG inhibitory neurons had substantially altered theta phase locking, with much more dispersed firing compared to healthy mice, particularly in males. Furthermore, optogenetically identified PV+ interneurons were specifically altered, while SOM+ interneurons could not be recorded, presumably due to cell death of this population in the DG. We therefore predicted that altered PV+ interneuron phase locking may directly contribute to seizures in epilepsy.

To test this hypothesis, we developed, validated, and utilized a new tool, PhaSER, to control the phase locking of PV+ interneurons unilaterally during behavior and tested their causal role in seizure susceptibility. We found that re-aligning PV+ interneuron firing to the trough of theta was able to reduce seizure susceptibility in male epileptic mice, while disrupting normal PV+ interneuron phase locking in healthy mice increased seizure susceptibility. Together, these data provide the first causal evidence that DG PV+ cell theta phase locking directly and powerfully influences excitatory-inhibitory homeostasis in the hippocampal circuit and can bidirectionally control seizure susceptibility. This finding is especially compelling because we were manipulating a relatively small population of inhibitory neurons (∼75% of PV+ cells in the dorsal DG expressed opsin, and manipulations were performed unilaterally), yet we saw robust changes in susceptibility to seize after systemic kainic acid.

Surprisingly, we found that altered inhibitory spike timing was much more dramatic in epileptic male mice than epileptic female mice, despite them having similar seizure frequencies. This suggests that the mechanisms of seizure susceptibility in male and female epileptic mice may be different. Here we found clear evidence that male epileptic mice have disrupted inhibitory phase locking in DG, and that restoring PV+ interneuron phase locking to the trough of theta reduces seizure susceptibility (Fig 4). However, in female epileptic mice, where inhibitory phase locking is less disrupted (Fig S2), we found that re-aligning PV+ interneuron activity had no effect (Fig S8A). This null result is similar to what we found in healthy animals that received trough stimulation (Fig 4C). In both cases, there was already intact inhibition near the trough of theta, and adding further inhibition at this phase had no effect on seizure susceptibility. Therefore, disrupted inhibitory phase locking is sufficient to induce seizure susceptibility, but is not always necessary for seizure development. Indeed, epilepsy is a complex and heterogeneous disorder, as illustrated by the many neurological and psychiatric disorders with comorbid seizures and the variability in treatment outcomes with anti-epileptic drugs. Thus, a personalized approach may be needed across patients to treat seizures.

Though theta phase locking has been widely observed and studied, interrogating its causal role in network function is not possible with traditional approaches. Technical advances in large-scale electrophysiological data collection and real-time signal processing have opened the door for closed-loop, real-time, phase-specific manipulations^3,58–63^. Here, we describe and validate PhaSER, our open-source, user-friendly platform that allows fast, flexible, and highly accurate delivery of perturbations at any specified phase of an oscillation. The PhaSER set up is unique because it estimates theta phase continuously (i.e., it does not solely detect maxima or minima), it triggers manipulations with outstanding accuracy, and when combined with artifact-free silicon probes it permits continuous monitoring of single-unit activity during stimulations. Here we demonstrated that PhaSER can precisely alter the theta phase locking of DG interneurons with excitatory and inhibitory stimulation patterns applied to target the theta trough or theta peak. Notably, trough and peak stimulation paradigms are virtually identical, with the same amount of red and blue light delivered, except that each is shifted by half a theta cycle (only ∼60ms). Despite this, these stimulation paradigms produced opposite effects on seizure susceptibility. This is powerful evidence that the timing of neural activity is a critical factor mediating behavior and that temporally agnostic stimulations may muddy experimental interpretability and produce mixed results in the clinic and at the bench. It is our hope that the open-source release of PhaSER will reduce hurdles to performing stimulations that are driven by the brain’s real-time endogenous activity.

Because seizures can serve as clear readouts of pathological network activity, the epileptic hippocampus is ideal for causal investigations of cell-circuit interactions such as theta phase locking. However, phase-locked spiking is also disrupted in models of substance use, schizophrenia, traumatic brain injury, autism, and Alzheimer’s disease^15–22^. Notably, all of these disorders also have increased risk for seizures^64–69^. This suggests that phase locking disruptions may be a relatively universal component of network dysfunction, including seizure susceptibility, across diseases. Therefore, our findings expand our fundamental understanding of inhibitory theta phase locking by directly demonstrating their causal role in seizure susceptibility, and inform the development and expansion of oscillation-driven neuromodulatory therapies aimed at restoring network function in epilepsy and beyond^70–75^.

## MATERIALS & METHODS

All experimental protocols were approved by the Icahn School of Medicine at Mount Sinai’s Institutional Animal Care and Use Committee, in accordance with the US National Institutes of Health guidelines.

### Animals

The SOM-Cre and PV-Cre mice used for experiments were bred and maintained in-house. SOM-Cre mice (Jackson Laboratory strain #013044; Sst^tm2.1^(cre)^Zjh^/J) were maintained homozygous^76^. The initial PV-Cre mice used for opto-tagging experiments (Fig 1) were maintained homozygous (Jackson Laboratory strain #017320; B6.129P2-Pvalb^tm1^(cre)^Arbr^/J)^77^. Later (Figs 3-4) we switched to maintaining this strain heterozygous by crossing male PV-Cre mice with female C57BL/6 (Charles River) mice to improve pilocarpine-induced status epilepticus success rates. For confirmation of viral specificity in mice without Cre expression, C57BL/6 (Charles River) mice were purchased directly from the vendor and allowed to habituate for at least a week before saline or pilocarpine treatment. Male and female mice were used for all experiments. Mouse sex was identified based on their external genitalia at the time of weaning (postnatal day 21). Data from male and female mice are combined except where noted. All mice were housed in standard housing conditions (12-hour light, 12-hour dark) in the animal facility at the Icahn School of Medicine at Mount Sinai. Animals were group-housed with their littermates when possible, or with an ovariectomized female (Jackson Laboratory strain# 000691; 129X1/SvJ) when littermates were not available or compatible. Animals were allowed *ad libitum* access to food and water except when water-restricted for training on the virtual linear track.

### Epileptogenesis with Pilocarpine

Chronic temporal lobe epilepsy was induced using the pilocarpine-status epilepticus (pilo) model as previously described^16,17^, with modifications to account for mouse strain differences. At ∼9 weeks of age, mice were injected with scopolamine methyl bromide (intraperitoneal, 1mg/kg, Sigma-Aldrich, Cat# S8502) to reduce the peripheral effects of pilocarpine. 30 minutes later, mice received intraperitoneal injections of pilocarpine hydrochloride (Sigma-Aldrich, Cat# P6503) to induce status epilepticus, or an equivalent volume of saline (Fresenius Medical Care, Cat# 060-10109) for control (i.e., healthy) mice. Initial pilocarpine dosage and booster protocols varied by strain and animal sex, each optimized to increase SE induction success rates. C57BL6 mice received initial pilocarpine dosages determined by their weight class (<20g mice received 285mg/kg, 20-25g mice received 275mg/kg, and >25g mice received 260mg/kg), with boosters of 50-100mg/kg delivered after 45 minutes if the animals had not seized. SOM-Cre mice received an initial pilocarpine dose of 365mg/kg for males and 345mg/kg for females, with boosters for both sexes (∼33% volume of initial dose) delivered 30-45 minutes after the initial dose as needed until the mouse began having seizures. Additional boosters were given if the time between seizures was greater than 10 minutes. Homozygous PV-Cre mice received an initial pilocarpine dose of 400mg/kg, with boosters (∼25% volume of initial dose) delivered 45-50 minutes after the initial dose as needed, with follow-up boosters every 20 minutes until SE was reached. Heterozygous PV-Cre mice received an initial pilocarpine dose of 285-315mg/kg, depending on experimenter, with boosters (∼25% volume) delivered 45-50 minutes after the initial dose as needed. We found that SE-success rates in male heterozygous PV-Cre mice were particularly sensitive to experimenter sex, with male mice ∼2.3x more likely to enter SE when a male experimenter was present (or when exposed to male experimenter scent via KimWipe or cage change the night before epileptogenesis).

SE was established when animals entered a continuous seizure and didn’t react to outside stimuli. Pilocarpine-treated mice were left in SE for 2 hours, after which they received diazepam (20mg/kg, i.p.; Dash Pharmaceuticals, NDC 69339-137-01) to terminate SE. Saline-treated animals were given equivalent injections of saline (rather than pilocarpine) and diazepam time-locked to the pilocarpine-treated mice. Pilocarpine-treated mice who didn’t reach SE were excluded from the study.

### Stereotaxic Surgery

The intrahippocampal virus delivery, headbar implant, and (where applicable) the wireless EEG implantation was performed during the same surgery under 1-3% isoflurane anesthesia. First, the skull was exposed and a small burr hole was drilled above the viral delivery site: 2mm posterior to bregma and 1.35mm to the right of bregma. A Nanoject III (Drummond Scientific Company) with glass capillary loaded with virus was lowered into the burr hole until the tip was 2.1mm ventral to bregma to target the DG. Virus was injected at a rate of 2nL/s. Once the injection was complete, the syringe was left in place for a minimum of 3 minutes. Then, for experiments where animals received viral injections into CA1 as well, the syringe was raised to 1.5mm ventral from bregma where an additional bolus of virus was injected in the same manner.

To determine baseline firing profiles of optogenetically-identified PV+ or SOM+ cells in the DG of control and epileptic mice, and monitor responses to theta phase-locked manipulations, a Cre-dependent channelrhodopsin virus (ChR2; AAV1-EF1a-double floxed-hChR2(H134R)-EYFP-WPRE-HGHpA; Addgene cat #20298; 1.2×10^13^ GC/mL; gifted by Karl Deisseroth) was injected into the dorsal hippocampus (120nL per site, DG and CA1). To test how theta phase locking of DG PV+ or SOM+ cells impacts seizure susceptibility, a Cre-dependent soma-targeted BiPOLES virus (somBiPOLES; AAV1-hSyn-DIO-somBiPOLES-mCerulean, AAV construct produced by Virovek from Addgene plasmid #154951, 2.0×10^13^vg/mL, 200nL)^47^ or Cre-dependent fluorophore-only control virus (GFP; AAV1-hSyn-DIO-eGFP, Addgene #50457, 2.2×10^13^ GC/mL, 200nL; gifted by Bryan Roth) was injected into the DG.

Following viral infusion, a stainless steel headbar was fixed onto the skull of the mouse. Lidocaine (2%, ∼0.03mL) was first injected subcutaneously over the skull, and the epaxial muscles along the neck were cleared from the skull’s surface. The mouse skull was thoroughly scored and then was stereotactically aligned to the headbar before the headbar was fixed near the surface of the skull with cyanoacrylate glue and dental cement (Lang Dental). Dental cement was built up to create a well around the exposed skull which was then filled with Kwik-Sil (World Precision Instruments). Once dried, the Kwik-Sil was covered with a final layer of dental cement. Meloxicam (5mg/kg) or Carprofen (5mg/kg) was administered subcutaneously during and for 2 days following the surgery together with a 7-day course of ampicillin (20mg/kg).

### Viral Validation

To confirm that the Cre-dependent viruses used in these studies would not express in the absence of Cre, control and epileptic C57BL6 mice were injected with Cre-dependent ChR2 (N=4 control, N=6 pilo), Cre-dependent somBiPOLES (N=3 control, N=3 pilo) or Cre-dependent fluorophore-only (N=3 control, N=3 pilo) viruses targeting dorsal DG and CA1. At least 20 days following viral injection, mice were deeply anesthetized with isoflurane (∼5%, Isospire), quickly decapitated, and their brains were dissected and dropped in 4% paraformaldehyde (Wako) and kept at 4°C for ∼24 hours. 50µm thick coronal sections spanning the entire hippocampus were prepared in 1× phosphate buffered saline (PBS; Fisher BP399) on a vibratome (Leica VT1000S). Every section containing hippocampus was mounted for fluorescent imaging with DAPI (SouthernBiotech DAPI Fluoromount-Gm, 0100-20) when compatible with viral expression (i.e., when visualizing the ChR-eYFP or eGFP signals), and mounted without DAPI (Invitrogen ProLong Gold P36930 or Vector Laboratories Vectashield H-1000) when the viral fluorophore was not compatible with DAPI (i.e., when visualizing the somBiPOLES-mCerulean virus). Every section was then viewed and imaged (Leica DM6B). No expression was noted in any of these animals, confirming that these Cre-dependent viruses do not express in the absence of Cre.

To validate Cre-dependent viral expression against immunohistochemistry, animals (N=3 PV-Cre control, N=3 SOM-Cre control, N=2 PV-Cre pilo, N=2 SOM-Cre pilo) were injected with AAV1-hSyn-DIO-somBiPOLES-mCerulean into the dorsal DG. 9-10 weeks after viral injection, they were deeply anesthetized with isoflurane before being perfused with ice-cold 1× PBS and 4% paraformaldehyde. Their brains were dissected and incubated in 4% paraformaldehyde at 4°C for <24 hours before being transferred to 30% sucrose in PBS. After brains were saturated with sucrose (∼2 days), they were frozen at -80°C until sectioning at 40µm on a cryostat (Leica CM3050S).

Immunohistochemistry was performed on every fourth section to stain for either somatostatin (SOM; rat anti-somatostatin, EMD Millipore MAB354, 1:300) or parvalbumin (PV; rabbit anti-parvalbumin, Swant PV27a, 1:1000) in red (goat anti-rat Alexa Fluor 555, Invitrogen A21434, 1:500 or goat anti-rabbit Alexa Fluor 568, Invitrogen A11036, 1:500, respectively). Sections were then mounted with Vectashield H-1000 and imaged on a confocal microscope (Zeiss LSM780). Colocalization of virus with immunofluorescence was determined by an experimenter blinded to genotype and experimental group. Manual counts of mCerulean-labeled cells were performed and comparing against PV or SOM immunofluorescence in three sections per mouse: anterior to the injection site (1.8mm posterior to bregma), at the injection site (2mm posterior to bregma), and posterior to the injection site (2.2mm posterior to bregma).

In addition, viral expression was verified alongside probe tract imaging for all recorded animals: virally expressing cell bodies were confirmed in the strata and cell morphologies we would expect (i.e., Cre-dependent viral expression in PV-Cre mice was primarily found in somata located on the hilar border of the granule cell layer, with dense axons throughout the granule cell layer, and expression in SOM-Cre mice was primarily found in somata located in the hilus, with axons densest in the molecular layer) and animals without viral expression were excluded from further analysis. No animals, control or epileptic, showed unexpected or inappropriate labeling.

### Chronic Seizure Monitoring

To chronically monitor seizures, a wireless EEG transmitter (PhysioTel ETA-F10; Data Sciences International) was implanted under the skin, posterior to the left ribcage, but anterior to the left haunch. The EEG transmitter leads were fixed between the skull and dura in small burr holes. The signal wire was fixed with cyanoacrylate glue under the left frontal bone, and the ground wire was fixed under the right side of the interparietal bone. EEG signals were monitored via Ponemah and NeuroScore software platforms (Data Sciences International), and seizures were manually identified. Pilocarpine-SE epileptogenesis produces highly stereotyped behavioral seizures that can be easily identified in the EEG signal by eye, and we have previously performed blinded video monitoring to confirm that these EEG seizures correspond to behavioral seizures ^16^. EEG data from N=10 control, saline-treated mice (combined genotypes) was assessed by a blinded experimenter, and no seizures were found. Therefore, to increase throughput in follow-up experiments (Fig 4, Fig S8), some control mice were implanted with a sham transmitter (ETA-F10 Training Module, Data Sciences International).

### Virtual Reality (VR) Training

In the days following surgery, mice were gently handled for 5 minutes daily. Once recovered from the surgery and fully habituated to being handled (∼3 days), mice were introduced to head-fixation (5 minutes per day for ∼3 days). Mice were then water restricted and maintained at a body weight of ∼82% of their initial weight while they continued the rest of their daily training (as in ^16^). During water restriction, mice were weighed and monitored daily for signs of dehydration, fatigue, or infection and were immediately given free access to water if any signs of distress were observed. After water restriction began, mice were introduced to head-fixation while standing/walking on a spherical treadmill locked to rotate on only one axis. Once the mice had gained coordination and strength on the spherical treadmill (usually ∼3 days of training for 10-20 minutes per day), mice were trained to lick from a water port that delivered a 4µL drop of water per lick. Once the mice had learned to reliably earn water from the water port, they were trained to run along virtual linear tracks of increasing length. For animals expressing ChR (included in Figs 1, S1, S2, S4, and S5), water rewards were delivered automatically at the end of the track, before the mice were teleported back to the beginning of the track. For animals expressing BiPOLES (included in Figs 2, 3, and S7), mice had to stop inside a reward zone to earn water^78^. Note that animals used for testing acute seizure susceptibility (Figures 4, S8) were not trained in VR. The virtual linear tracks were created with ViRMEn^79^ and were displayed across three flat monitors angled around the front of the spherical treadmill. Once the mice were consistently earning their daily water within 1 hour on the longest (∼2 meter) virtual track, they were prepared for acute silicon probe recording. Virtual track training took ∼5 days.

### Craniotomy & Ground Implantation Surgery

The day prior to an acute silicon probe recording, a craniotomy and ground implantation was performed. First, the most superficial layer of dental cement was drilled off and the Kwik-Sil was removed, exposing the skull. Then a burr hole was drilled above the left hemisphere of the cerebellum, and an Ag/AgCl-coated silver reference wire (Warner Instruments) was slipped between the skull and dura and fixed with cyanoacrylate glue and dental cement. A 1.5mm diameter craniotomy was also drilled over the right hippocampus at this time (centered on the following coordinates: 2mm posterior, 1.45mm right from bregma). The craniotomy was covered with buffered artificial cerebrospinal fluid (ACSF; in mM: 135 NaCl, 5 KCl, 5 HEPES, 2.4 CaCl2, 2.1 MgCl2, pH 7.4)^16,80^ and the exposed skull was again covered with Kwik-Sil. Mice were returned to their home cages overnight after recovering on a heating pad.

### Acute Silicon Probe Recordings & Recording Hardware

The day following craniotomy, mice were set up for acute silicon probe recordings [15.5 ± 0.5 weeks of age (109 ± 4 days of age; 19 ± 1 day after initial surgery)]. First, a 64 channel Cambridge NeuroTech (ASSY-77-H3) silicon probe (single shank, 8mm length, sharpened tip) with attached lambda-b optic fiber 100µm core (tapered 1.2mm) was painted with dye (Invitrogen, Vybrant DiI, V22885) so the probe track could be visualized later. Mice were then head-fixed on the spherical treadmill and the Kwik-Sil covering the skull was replaced with a buffered ACSF solution. Head-flat skull positioning in the dorsal-ventral and medial-lateral directions was then established to within 50µm (bregma-to-lambda) before the silicon probe was lowered into the right dorsal hippocampus at the following approximate coordinates: 2mm posterior, 1.45mm lateral, 2.1mm ventral to bregma. After reaching the target depth and confirming probe location with electrophysiological signatures of the dorsal hippocampus, mineral oil was placed over the buffered ACSF solution and the probe was allowed to settle for 1 hour before starting the virtual linear track and recording^16,81^.

Electrophysiological signals were sent to an Intan headstage (RHD 64-Channel Recording Headstage, Intan Technologies) via a Samtec to Omnetics adaptor (ADPT A64-Om32×2, Cambridge NeuroTech) for pre-amplification and digitization. An Intan recording controller (RHD2000 Intan 1024ch Recording Controller, Intan Technologies) collected and logged signals at a 25kHz sampling rate from each electrode channel as well as from analog inputs reflecting the speed of the spherical treadmill, position on the virtual linear track, time of water delivery, time of the mouse’s licking behavior, and LED trigger and delivery times. Because the Intan recording controller logs data to the PC over a USB connection, which is relatively slow, we selected a single electrode channel in the deep CA1 pyramidal cell layer (as identified by its location relative to a stereotypical theta phase shift and higher density of spikes) to route as an analog output going to a separate, low-latency, PCIe data acquisition device (NI-DAQ PCIe-6321, National Instruments).

For recordings on mice expressing ChR2 (Figs 1, S1, S2, S4, S5): mice ran down the virtual linear track at least 50 times to collect a baseline before any photo-manipulations were introduced. Immediately following this baseline period, single-units were photo-tagged with 100 1Hz square pulses (10ms duration) of blue light (LED, 455nm, Prizmatix, 1mW) sent via a Pulser Plus (Prizmatix) pulse train generator. Five minutes following the photo-tagging procedure, a baseline of at least 120s was acquired to train the auto-regressive model for phase estimation (described in detail below). Photo-stimulation locked near the trough (170°) or peak (0°) was then delivered for at least 25 trials during locomotion. No phase-specific photo-stimulations were delivered during rest. The order of trough- and peak-targeted stimulations was counterbalanced across animals with at least 5 minutes between the two manipulation periods.

For VR recordings on mice expressing BiPOLES (Figs 2, 3, S7): mice ran down the virtual linear track 25 times to collect an initial baseline before any photo-manipulations were introduced. Then, trough (170°) or peak (0°) targeted red light stimulations (10ms; LED, 625nm, Prizmatix, <0.72mW), with blue inhibitory light pulses (variable duration) targeted ±90° around the opposite phase (from 270° to 90°, or from 90° to 270°, respectively), were delivered in 25 trial intervals with baseline periods in between. Trough and peak targeted manipulations were counterbalanced across animals (e.g., 25 trials baseline, 25 trials trough/peak, 25 trials baseline, 25 trials peak/trough, 25 trials baseline). These manipulations were delivered regardless of locomotion status, but single-unit and LFP analyses were limited to periods when the mice were running.

### Testing Seizure Susceptibility with Kainic Acid

To test whether PV+ or SOM+ cell theta phase locking impacted seizure susceptibility, control or epileptic PV-Cre or SOM-Cre animals were randomly assigned to receive an injection of either an opsin-(AAV1-hSyn-DIO-eGFP) or opsin+ virus (AAV1-hSyn-DIO-somBiPOLES-mCerulean) in the dorsal DG. At the same time, mice underwent surgery to implant a headbar and a wireless EEG telemetry device (or sham device for some control animals) to monitor spontaneously occurring seizures. Following recovery from surgery, mice were habituated to experimenters with gentle handling for 3 days (5 minutes per day). Once habituated to the experimenters, mice were then habituated to head-fixation for an additional 3 days. At this point, mice were introduced to the Styrofoam ball used for head-fixed recordings and trained to walk on the ball for increasing durations until comfortable (minimum of 3 days). Simultaneously, mice were habituated to receiving intraperitoneal injections of 0.2mL saline while head-fixed on the ball. Once fully habituated to the experimental set-up, mice underwent craniotomy and ground implantation surgery, and the seizure susceptibility test was performed the following day. Details of the craniotomy and recording hardware are described above. Note that in a few cases, the seizure susceptibility recording was delayed by a day or two if spontaneous seizures occurred within 4 hours of the planned silicon probe recording.

To test if PV+ or SOM+ cell phase locking impacts seizure susceptibility, an acute silicon probe was lowered into the right dorsal hippocampus, with channels spanning CA1 and the DG, of control or epileptic PV-Cre or SOM-Cre mice expressing either eGFP (opsin-) or BiPOLES (opsin+) virus. Mice were recorded for 30 minutes to establish a baseline and ∼2 minutes of baseline data from a channel in CA1 deep pyramidal cell layer was used to train PhaSER’s auto-regressor (details below). Mice were then injected intraperitoneally with 2mg/mL kainic acid (KA; Tocris Bioscience, 7065) mixed in saline (30mg/kg KA for control animals; 20mg/kg for epileptic animals). Immediately following KA injection, phase-specific red/blue light delivery began with excitatory red light pulses (10ms) targeted at either the peak (0°) or trough (170°) of CA1 theta, and inhibitory blue light pulses (variable duration) targeted ±90° around the opposite phase (from 90° to 270°, or from 270° to 90°, respectively). Light delivery continued until the experimenter visually confirmed electrographic seizure onset. Shortly after seizure cessation, the recording was ended and the mice were swiftly euthanized and their brains dissected for histology.

Animals were excluded from analyses if the mean phase-angle of excitatory (red) light delivery was >45° from the target phase (N=10 of 121 animals), or if the excitatory red light deliveries were not phase-specific (N=1 animal, Rayleigh test for non-uniformity of circular data). Other reasons for exclusion included poor KA injection, no seizure within 70 minutes of KA injection, seizure occurrence in the 4 hours leading up to KA injection, lack of viral expression, mis-targeted silicon probe, severe hippocampal sclerosis that prevented CA1 theta referencing, or evidence of seizure-like activity occurring before the KA injection in a control mouse.

### Real-time Signal Processing & Phase-Specific Manipulation with PhaSER

In order to apply light stimulations locked to the ongoing theta oscillation in real-time, we created a custom LabVIEW program, which can be found on our lab’s GitHub (https://github.com/ShumanLab) along with user-friendly documentation. The 25kHz recorded signal from the CA1 pyramidal layer reference channel was first down-sampled to 1kHz. Then, in order to do theta phase detection, we filtered the down-sampled signal for the theta band (4-9Hz) in real-time. Because digital filters introduce phase lags and distortions, a zero-phase filter (first-order Butterworth) was employed. Zero-phase filtering is a technique in which a digital filter (in this case, an infinite impulse response (IIR) filter), which introduces a phase lag, is applied first in the forward direction and then in the reverse direction to cancel out the phase lags in the output signal. To extract the theta phase from this filtered signal, we used the Hilbert transform. The Hilbert transform can be used to calculate the phase angle of a real signal at every point in time, however both digital filters and the Hilbert transform introduce edge effects – distortions at the edge of the analysis window. To circumvent these distortions, and therefore avoid incorrectly identifying the theta phase, we removed 150 samples from the beginning and end of the filtered signal within the 1000 sample sliding filter window and then used a 13th order auto-regressive model that has been described previously^46^. The auto-regressive model was trained on ∼2 minutes of baseline data using the Yule-Walker method. This model was then used to reconstruct the 150 most recent datapoints without edge effects and then forward predict an additional 150 samples (which equals approximately 1 theta cycle). This places the current time point at the center of the 300-sample prediction window (Fig S3). The Hilbert transform was then applied to the resulting signal and the current phase angle was calculated from the center of the phase-estimation window, which corresponds to the real-time signal without edge effects. A schematic of this process is shown in Fig S3. To limit photo-stimulation to periods when the animal was locomoting, a movement threshold was set based on the analog signal from the spherical treadmill. When this value passed the threshold indicating locomotion, and the Hilbert transform phase angle crossed the threshold of the light delivery target phase, a TTL trigger was generated by the NI-DAQ and passed to a pulse generator (Pulser Plus, Prizmatix) which was programmed to deliver 10ms of blue or orange-red light (LED, 455nm or 625nm, Prizmatix, ∼1.7mW blue light power, ∼0.72mW red light power) at the target phase. The theta trough stimulation target was 170° and the theta peak stimulation target was 0°. When bidirectional manipulations were applied, a TTL step was generated by the NI-DAQ to toggle the blue LED on 90° after the target phase and off 90° before the following target phase (i.e., inhibitory blue light was applied ±90° around the phase opposite the stimulation target phase). The latency of this real-time signal processing system from electrode to light trigger is approximately 3ms, with the majority of the latency produced when the data is read from the DAQ.

### Histology Following Recordings

Following recording, the silicon probe was removed from the brain and the mice were deeply anesthetized with isoflurane (5%; Isospire) before being decapitated. The brains were quickly dissected and dropped in 4% paraformaldehyde (Wako) where they stayed for ∼24 hours. 50µm thick coronal sections were prepared in 1× phosphate buffered saline (PBS; Fisher BP399) using a vibratome (Leica VT1000S). All sections containing hippocampus were then mounted for fluorescent imaging with DAPI (SouthernBiotech DAPI Fluoromount-Gm, 0100-20) when compatible with viral expression (i.e., when visualizing the ChR-eYFP signal), and mounted without DAPI (Invitrogen ProLong Gold P36930; or Vector Laboratories VECTASHIELD H-1000) when the viral fluorophore was not compatible with DAPI (i.e., when visualizing the somBiPOLES-mCerulean virus). Sections were later imaged to confirm viral expression (visualized via eYFP or mCerulean) and to determine probe location (DiI). Images were taken using a Leica DM6B fluorescence microscope equipped with a Lumencor Light Engine and Leica DFC9000 GT camera. Hippocampal anatomical locations of the probe tracks and stereotaxic coordinates were compared to their electrophysiological signals and a mouse brain atlas^82^. Viral expression was confirmed and compared to expected expression patterns (e.g., expected expression of Cre-dependent virus in SOM-Cre mice was in CA1 stratum oriens and DG hilus while expected expression in PV-Cre mice was in CA1 pyramidal cell layer and on the border of DG hilus/granule cell layer) in all recorded mice.

#### Post-Processing & Analysis of Single-Unit and Local Field Potential Data

All data analysis was performed with custom scripts using MATLAB 2017a. Data files were first concatenated into continuous signals for each channel. To extract single-units, all channels were high-pass filtered and background subtracted for clustering into putative single-units using Kilosort2.5^83,84^ and Phy2^84^. This automated spike sorting pipeline reliably isolates single-units using a template-matching approach that is iteratively updated and includes drift correction, a critical step for stable cell isolation across the entire recording. Once single-units were isolated using Kilosort2.5 and manually confirmed in Phy2, we calculated the percentage of spikes in each putative cell cluster that violated a 2ms refractory period, and only continued analysis with clusters where 3% or fewer spikes violated this refractory window. We then assigned cell clusters as “inhibitory,” “excitatory,” or “unclassified” based on their firing rate, distribution of inter-spike intervals, and complex spike index (Fig S1A,D). We then tested each single-unit for its response to our photo-tagging stimulation protocol. To identify ChR2-expressing cells, we used a 1Hz 10ms blue light tagging protocol and built a spike probability distribution based on the 500ms prior to each of 100 light stimulations. If a single-unit’s spike frequency during the first 4ms of light delivery was significantly above the distribution mean (where α = 0.001), then the cell was classified as ChR2+. To identify BiPOLES-expressing cells, we used a similar 1Hz photo-tagging protocol, but tested for bidirectional responsiveness to 100 excitatory 10ms red light pulses and 100 inhibitory 20ms blue light pulses. To be identified as BiPOLES+, cells had to be both excited by red light and inhibited by blue light. A cell was classified as ‘excited by red light’ if the spike probability was significantly higher during the 10ms red light delivery than during the 500ms leading up to each light stimulus (α = 0.05), and was classified as ‘inhibited by blue light’ if the spike probability during the 20ms of blue light delivery was significantly lower than the 500ms prior (α = 0.05), or if there were fewer spikes in the 20ms blue light stimulation period than in any of the equivalently sized bins in the 500ms prior. Due to our histological confirmation that neurons labeled with our Cre-dependent viruses in healthy and epileptic PV-Cre mice co-localized with PV immunofluorescence, and those labeled in healthy and epileptic SOM-Cre mice co-localized with SOM immunofluorescence (and did not co-localize with PV immunofluorescence), we will refer to opto-tagged cells in the DG of PV-Cre mice as PV+ cells and opto-tagged cells in the DG of SOM-Cre mice as SOM+ cells. Putative inhibitory cells and opsin-expressing PV+ and SOM+ cells in the DG were then assessed for their phase preference during the baseline period as well as their response to the trough and peak phase-specific manipulations. Phase locking was quantified by an r-value (magnitude of phase preference) and a mu-value (circular mean phase of firing) relative to the deepest pyramidal layer channel (i.e., the pyramidal cell layer channel closest to the stratum oriens). Note that mu-values are only reported for cells that were significantly phase-locked during baseline (Rayleigh’s test for non-uniformity), whereas r-values are reported for cells regardless of the strength of their phase preference.

For local field potential (LFP) analyses, signals were down-sampled to 1kHz, bandpass filtered (theta: 5-12Hz) using zero-phase digital filtering and LFP power was compared using the Chronux library^85,86^. Power spectral density (PSD) was calculated from the unfiltered, down-sampled LFP at each electrode position during running bouts that were at least 3 seconds in duration. A spectrogram was constructed by taking the mean PSD across running bouts within each frequency bin (1Hz bin size) and electrode channel. Sublayers of the hippocampus were identified from histology showing location of the dye-coated probe and electrophysiological markers, including peak theta and ripple power, theta phase shifts, gamma coherence, and density of action potentials^87–90^. Channels from each layer were aligned and spectrograms for each stimulation condition were averaged across animals (Fig S7).

To determine if stimulating SOM+ or PV+ neurons reset the theta phase, we aligned the down-sampled LFP from the deepest CA1 pyramidal cell layer channel to the onset of red light pulses (10ms duration) delivered at 1Hz. The LFP was averaged across 100 light deliveries and the mean envelope was calculated during a 50ms wide bin starting 250ms before and after the aligned light deliveries. With the exception of the opto-tagging identification and theta phase reset analyses, during which the mice were spontaneously running and resting, all other single-unit and LFP power analyses were limited to periods when the animals were running.

#### Automated Detection of KA Injection, Seizure Onset, and Seizure Duration

Seizure latency was quantified as the amount of time between KA injection and seizure start time using data from one channel at the DG hilus/upper granule cell layer border as a reference. The time of injection, seizure start, and seizure end were identified with custom MATLAB scripts, which were compared against recording notes and confirmed via manual inspection by a blinded experimenter.

To identify the time of KA injection, we detected the end of the signal artifact that was introduced by touching the mouse. To do this, we filtered the DG reference channel for 57-63 Hz and then calculated line length values over a 1 second sliding window for the recording duration. The line length values from the baseline period (first 30 minutes) established a threshold using a method similar to prior work^91^. Briefly, the threshold was initially defined as 2 standard deviations above the median baseline line length. Each baseline line length value was then compared to this threshold, and a “hit” was counted if the value met or exceeded the threshold. If any hits were recorded, the number of standard deviations (SDfactor) was increased by 0.5 and hit testing performed again. This process was repeated until there were no hits in the baseline period, and the injection detection threshold was subsequently set to that SDfactor. To detect the injection, we compared the line length values in the post-baseline injection time window to this threshold. If no threshold crossings were detected, the SDfactor was decreased by 0.5 and the test was repeated again. The last threshold crossing was considered the injection time, corresponding to the last time in the injection window when 60Hz noise was higher than baseline.

The time of seizure onset (used to determine seizure latency from KA injection) was detected using a similar hit testing strategy, however line lengths were calculated from unfiltered LFP data from the DG reference channel and a moving average filter was applied to prevent short, non-seizure events (e.g., interictal spikes) from triggering seizure detection. For the majority of recordings (89 of 118), the baseline period was used to set the crossing threshold. However, in about 25% of cases (29 of 118), LFP dynamics after the injection were considerably altered compared to baseline activity, causing false seizure onset detection times. These changes in the LFP can be largely attributed to changes in movement (i.e., some mice changed their locomotion status in response to the injection). In these cases, a variable duration “post-injection/pre-seizure” period (10 seconds after the detected injection end time to 60 seconds before the manually recorded seizure start time) was used to set the detection threshold instead. Seizure onset was then identified as the first upwards threshold crossing following the injection end time. No significant differences in seizure latency were found between opsin-animals of the same genotype and experimental condition that received either peak- or trough-targeted stimulation (unpaired t-tests, p>0.05), and therefore these stimulation paradigms were collapsed into their respective PV-Cre Control/PV-Cre Epileptic/SOM-Cre Control/Som-Cre Epileptic Opsin-groups for follow-up analyses.

The end of the seizure (used to calculate seizure duration), was identified as the first time following seizure onset when the signal line length dropped and stayed below 66% of threshold for a minimum of 5 seconds. In six cases, manual inspection by a blinded experimenter did not agree with the detected seizure end. In five of these six cases, the percent reduction was set manually at either 33%, 50%, or 90% threshold. In the remaining case, no seizure duration was calculated as the signal suggested the mouse continuously seized until the end of the recording.

When latency to seizure or seizure duration was normally distributed (Shapiro-Wilk test, *p*>0.05) and standard deviations were not significantly different between trough/peak/opsin-groups (Brown-Forsythe test, *p*>0.05), ordinary one-way ANOVAs were used to establish whether there was a significant difference between the stimulation paradigm (trough/peak/opsin-) means, with post hoc Dunnett’s multiple comparisons tests where appropriate. However, if latency to seizure or seizure duration was not normally distributed, a Kruskal-Wallis ANOVA was performed, with post hoc Dunn’s multiple comparisons tests where appropriate.

### Statistics

All statistics were performed in MATLAB 2017a or GraphPad Prism. For circular data, all phases were converted to radians and were analyzed using the Circular Statistics Toolbox in MATLAB^92^. Linear data is presented as mean ± standard error of the mean while circular data is presented as circular mean ± standard deviation. For comparisons of mean preferred phase preference (mu), Watson-Williams and circular k-tests were performed post-hoc to test for a change in mean phase preferences or concentration of phase preferences only if the Kuiper test showed a significant difference. Furthermore, only neurons that showed significant phase modulation (Rayleigh’s test for non-uniformity) were included in mu comparisons. Threshold for significance (α) was set to 0.05 unless otherwise noted. n=cells, N=mice. In figures * indicates p<0.05, ** indicates p<0.01, and *** indicates p<0.001 unless otherwise noted.

## SUPPLEMENTAL FIGURES

**Fig. S1.**
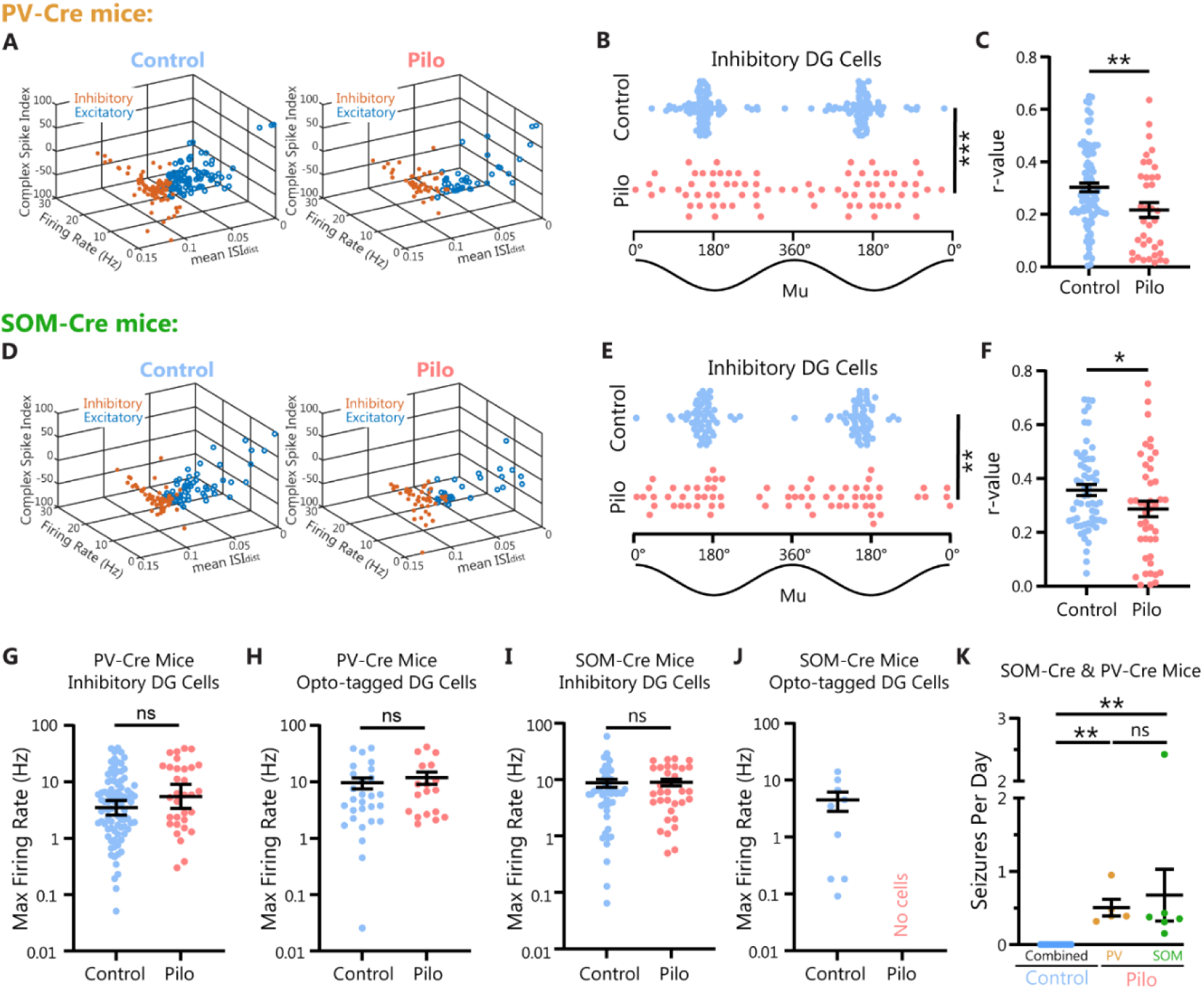
Disrupted DG interneuron theta phase locking in epileptic PV-Cre and SOM-Cre mice. **A-F)** Putative inhibitory DG cells from PV-Cre mice (A-C) or SOM-Cre mice (D-F). **A, D)** Spikes were clustered using Kilosort2.5 and then manually curated with Phy. Putative inhibitory and excitatory neurons in the dentate gyrus (DG) were distinguished by their spiking properties in control and epileptic PV-Cre (A) or SOM-Cre (D) mice. **B, E)** Inhibitory DG cells in epileptic PV-Cre (B) or SOM-Cre (E) mice show significantly altered theta phase locking profiles compared to controls [PV-Cre mice (B): Kuiper test p≤0.001, circular k-test p<0.0001, Watson-Williams test p=0.013; SOM-Cre mice (E): Kuiper test p≤0.001, circular k-test<0.0001, Watson-Williams test p=0.028]. Note that only cells that were significantly phase-locked to theta are included: PV-Cre (B): Pilo n=37 cells, N=4 animals; Control n=80 cells, N=8 animals; SOM-Cre (E): Pilo n=35 cells, N=5 animals; Control n=57 cells, N=6 animals. **C, F)** Inhibitory DG cell firing in epileptic PV-Cre (C) or SOM-Cre (F) mice was significantly less modulated by theta phase (reduced r-values) than in control mice [PV-Cre (C): unpaired t test p=0.008 (Pilo n=51 cells, N=5 animals; Control n=88 cells, N=8 animals); SOM-Cre (F): unpaired t-test p=0.043 (Pilo n=44 cells, N=5 animals; Control n=59 cells, N=6 animals)]. **G-J)** Maximum firing rates (from theta-binned firing rates, 20° bins) were not different in inhibitory DG cells or opto-tagged PV+ cells between control and epileptic mice (G: inhibitory DG cells from PV-Cre mice, unpaired t-test p=0.068; H: opto-tagged DG PV+ cells, unpaired t-test p=0.528; I: inhibitory DG cells from SOM-Cre mice, unpaired t-test p=0.896; J: opto-tagged DG SOM+ cells, note that no opto-tagged SOM+ cells were identified in the DG of epileptic SOM-Cre mice). **K)** Pilocarpine-induced SE produced chronic spontaneous seizures in PV-Cre and SOM-Cre mice, while saline-treated controls (combined genotypes) do not seize (Kruskal-Wallis test p<0.0001, Dunn’s posthoc tests Control vs PV-Cre Pilo adjusted p=0.001, Control vs SOM-Cre Pilo adjusted p=0.004, PV-Cre Pilo vs SOM-Cre Pilo adjusted p>0.999, N=10 Control, N=5 PV-Cre Pilo, N=6 SOM-Cre Pilo mice).

**Fig. S2.**
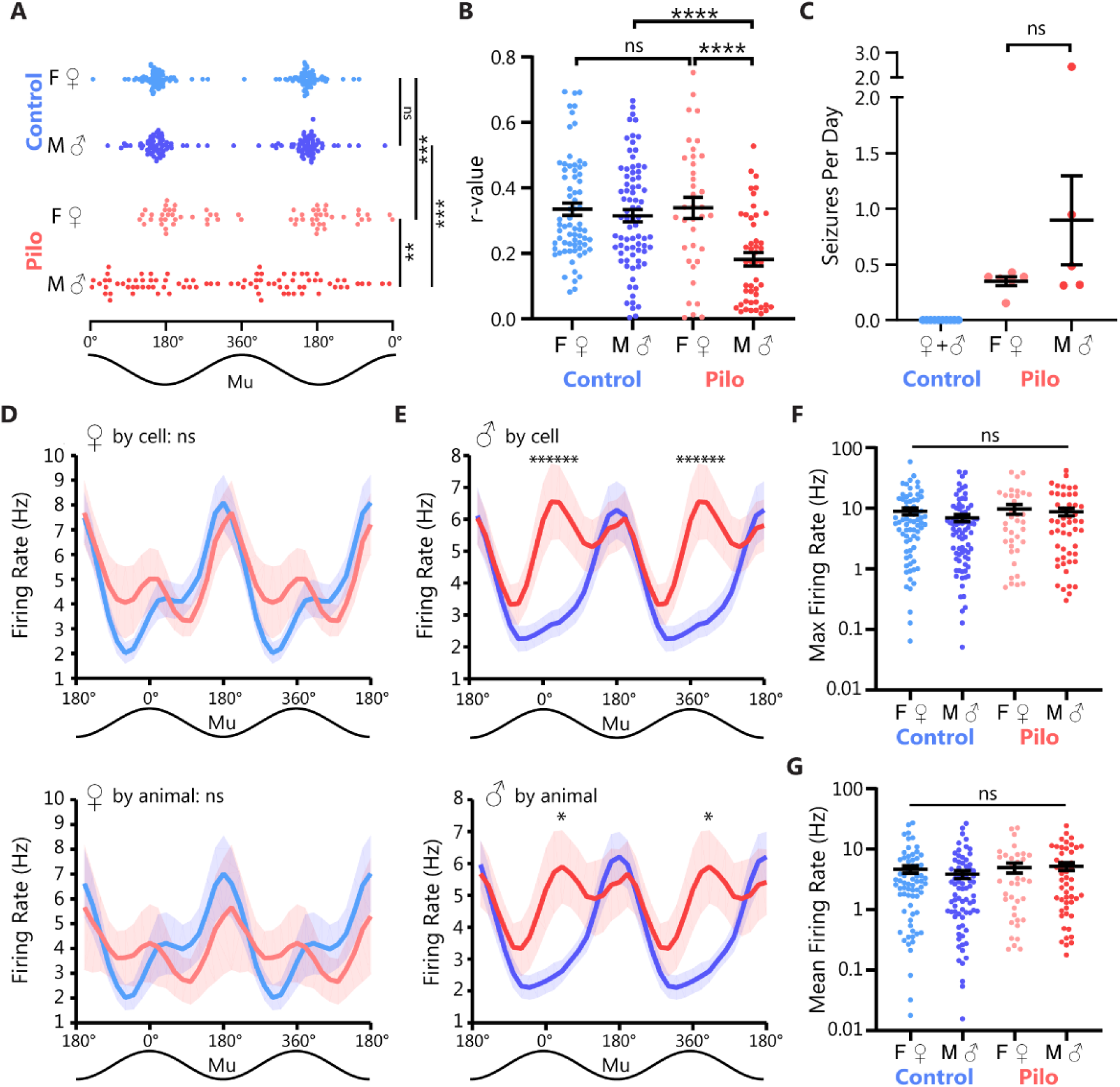
DG inhibitory theta phase locking is severely disrupted in epileptic male mice. **A)** DG interneurons from epileptic male mice showed a uniform distribution of mu values (mean theta phase preferences; Rao’s test for non-uniformity p=0.50), whereas DG interneurons in epileptic females and control animals of both sexes have phase preferences coupled to the trough of theta (Rao’s test for non-uniformity, p<0.001 for each group; PV-Cre and SOM-Cre genotypes combined). There are sex-specific alterations in DG inhibitory theta phase locking in epileptic mice (Kuiper test, male control vs male pilo p≤0.001, female control vs female pilo p=0.001, female control vs male control p=1.0, female pilo vs male pilo p=0.01). See Table S1 for summary of p-values when comparing mean phase preferences (Watson-Williams test) and concentration of phase preferences (k-test). **B)** DG inhibitory cells showed significantly diminished r-values (i.e., strength of theta modulation) in epileptic males compared to control males or epileptic females. DG interneurons in epileptic females showed no deficit in the strength of theta phase modulation compared to control females. Ordinary one-way ANOVA p<0.05 with Tukey’s post hoc comparisons shown. **C)** Pilocarpine epileptogenesis was effective at producing spontaneous, recurrent seizures in both male and female mice, with no difference in mean seizure frequency (unpaired Welch’s t-test p=0.242), though there was more variability in the epileptic males than females (F-test to compare variances p=0.0002). **D)** DG inhibitory cells’ firing rates fluctuated across the theta cycle (20° bins), with firing rates highest near the theta trough in control and epileptic females [top: by cell, two-way repeated measures ANOVA main effect of theta bin p<0.0001, of group (pilo vs control) p=0.784, interaction effect p=0.052; bottom: by animal, two-way repeated measures ANOVA main effect of theta bin p=0.021, of group (pilo vs control) p=0.213, interaction effect p=0.851]. N=7 control mice (n=70 cells), N=4 pilo mice (n=36 cells). Note that the theta cycle is double plotted for clarity. **E)** DG inhibitory cells’ firing relative to theta was dramatically altered in epileptic males, with the maximum firing rate shifting towards the peak of theta compared to control males (top: by cell, two-way repeated measures ANOVA main effect of theta bin p<0.0001, of group p=0.145, interaction effect p<0.0001, with posthoc tests showing significantly different firing rates from -20° to +100°; bottom: by animal, two-way repeated measures ANOVA main effect of theta bin p<0.0001, of group p=0.278, interaction effect p=0.003 with posthoc tests showing significantly different firing rates from 40° to 60°). N=5 control mice (n=77 cells), N=5 pilo mice (n=51 cells). **F)** No change in maximum firing rate (calculated from theta binned firing, as in D & E) between control and epileptic male and female mice (ordinary one-way ANOVA p=0.427). **G)** No change in mean firing rate (mean of theta binned firing, as in D & E) between control and epileptic male and female mice (ordinary one-way ANOVA p=0.505).

**Fig. S3.**
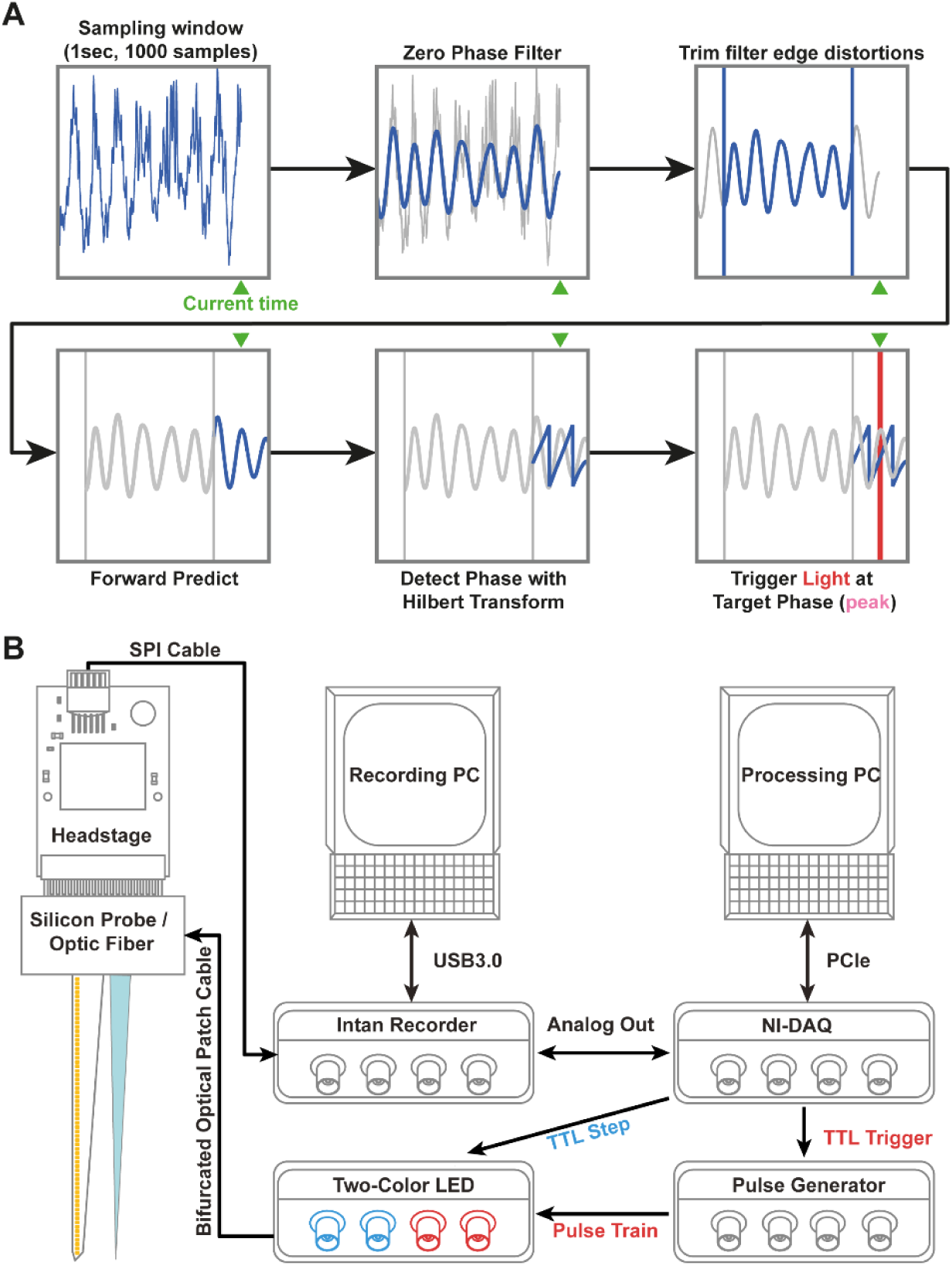
High-speed hardware and an autoregressive model allow PhaSER to predict and detect the theta phase in near real-time. **A)** A 1-second window of data is down-sampled to 1000Hz and is then zero-phase filtered and the first and last 150ms of the filter window are trimmed to remove edge distortions. An autoregressive forward prediction model is then used to extrapolate the trimmed 150ms followed by an additional 150ms forward prediction. Finally, the Hilbert transform is applied and the current phase is taken at the center of the 300ms forward prediction window (i.e., the current time). When the estimated current phase matches the target phase (in this example, the theta peak), excitatory red light delivery is triggered. The same process is used to estimate ± 90° from the target phase to toggle the inhibitory blue light on and off. **B) Electrophysiological data is digitized and amplified at the headstage before being passed into an Intan Recording System** and saved to a Recording PC. The theta reference channel is passed from the Intan Recorder as an Analog signal to a National Instruments DAQ (NI-DAQ). The signal is transferred to a processing PC via high-speed PCIe connector before undergoing phase estimation as described in A. Upon detection of the stimulation target phase, the NI-DAQ sends a TTL trigger to a pulse generator, which switches on a red LED connected via bifurcated optical patch cable to the implanted tapered optic fiber. Upon detection of the start and end of the inhibitory window, the NI-DAQ separately sends a TTL signal to directly toggle on (and then off) a blue LED connected to the same bifurcated optical patch cable. The electrophysiological signal is passed from the silicon probe to the processing PC using high speed SPI and PCIe cables to achieve low-latency (∼3ms) signal processing. Additionally, to prevent overburdening the CPU, we use separate PCs for recording and real-time signal processing, though this may not be strictly necessary.

**Fig. S4:**
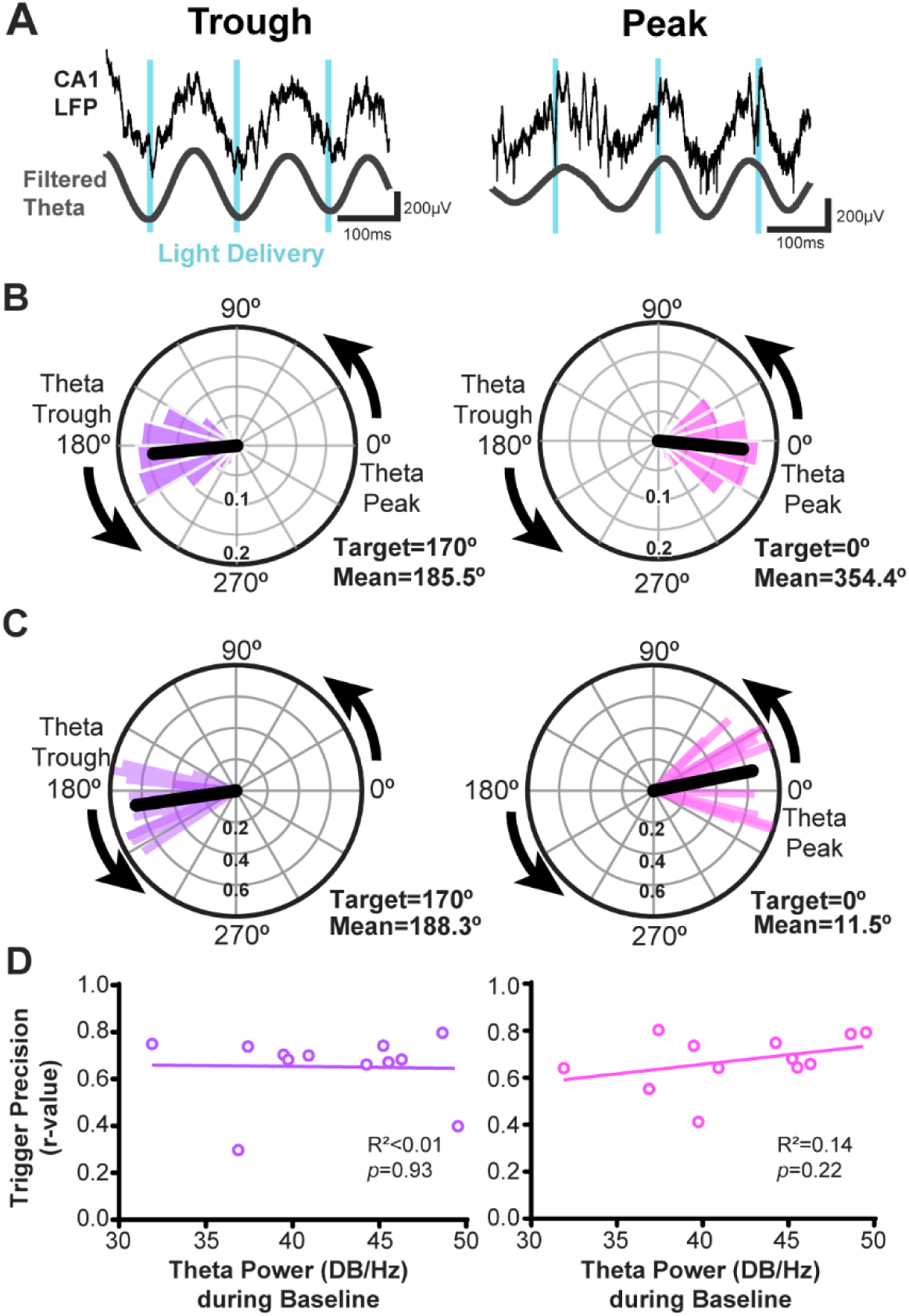
PhaSER accurately detects and triggers light delivery to a specified theta phase. **A)** Light delivery (blue) was delivered at the trough (left) or peak (right) of CA1 theta (both the raw reference LFP and offline filtered signal are shown). **B)** Circular distribution of light pulses in a representative animal targeted to the theta trough (left, target 170°) were centered around the theta trough (mean phase of light delivery = 185.5°) and light pulses targeted to the theta peak (right, 0°) were centered around the theta peak (mean phase of light delivery = 354.4°, or -5.6°). The mean phase is shown in black. **C)** Polar plot showing the distribution of the mean phase of light delivery across mice when targeting the trough (left, target 170°, mean = 188.3°) and peak (right, target 0°, mean = 11.5°) of CA1 theta. The trigger precision (r-value, length of the resultant vector) is shown on the radial axis. **D)** The trigger precision (r-value) targeted towards the trough (left) or peak (right) was not significantly correlated with theta power during baseline (trough: Pearson r=-0.03, p=0.93; peak: Pearson r=0.38, p=0.22, N=12 animals).

**Fig. S5:**
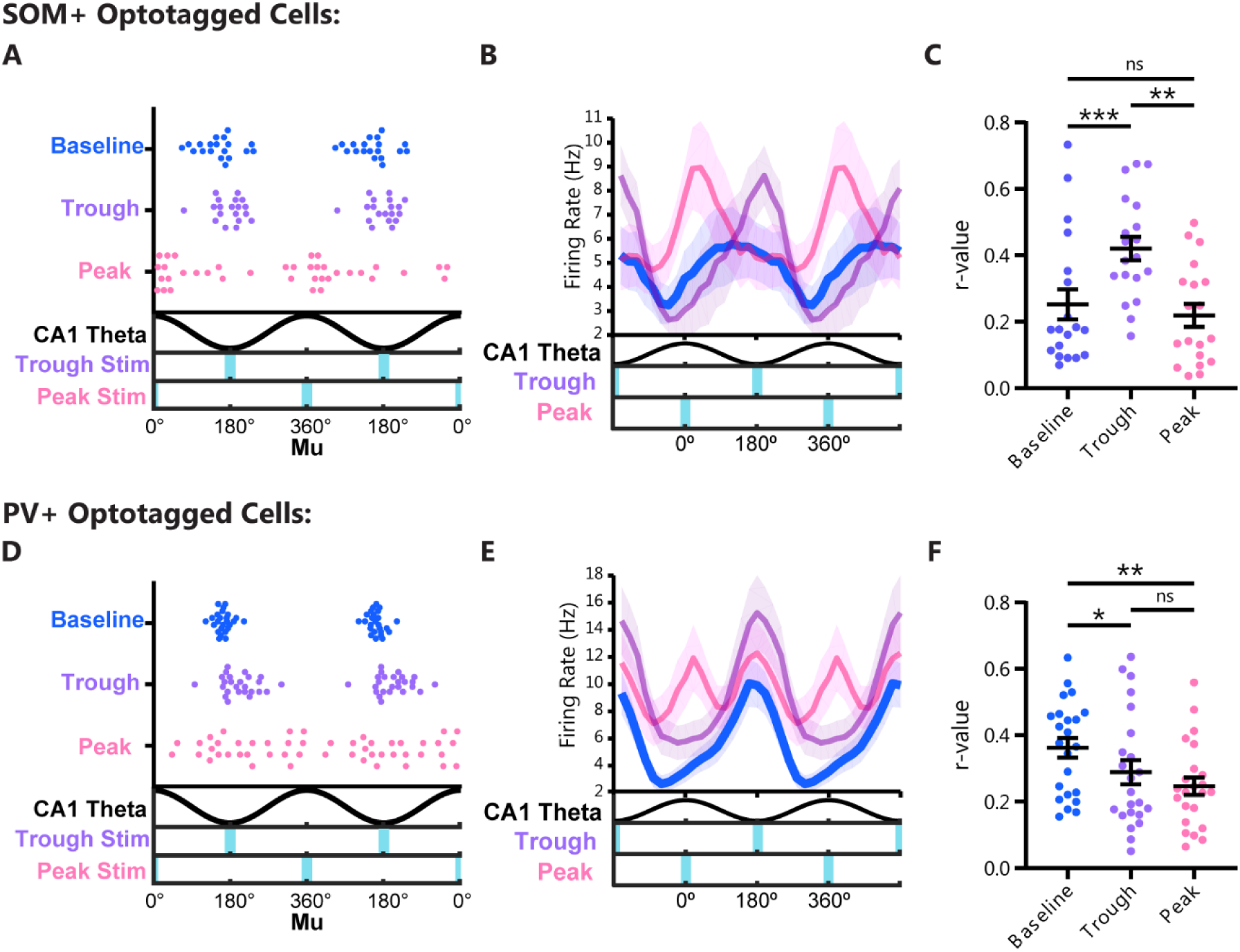
Phase-locked excitation in ChR2-expressing SOM+ or PV+ cells. **A)** Mean preferred firing phase (mu) of ChR2-tagged SOM+ neurons in DG and CA1 of healthy SOM-Cre mice during baseline (blue), trough-targeted stimulation (purple), and peak-targeted stimulation (pink; N=5 mice, n=19 cells). Trough-targeted stimulation did not significantly alter SOM+ cell theta phase preference (Kuiper test p>0.1), though it did bring the numerical mean phase preference closer to the true trough (baseline circular mean mu: 152.4°, with trough stimulation: 180.0°). Peak-targeted stimulation significantly shifted the mean preferred firing phase compared to baseline (32.2°; Kuiper test p≤0.002; Watson-Williams test p<0.0001), without altering the distribution of phase preferences compared to baseline (circular k-test p=0.21). **B)** Peak and trough-targeted stimulations increased SOM+ cell firing rates at the stimulated phase of theta (20° bins). **C)** The strength of phase preference, or r-value, was significantly increased during trough-targeted stimulation compared to baseline or peak-targeted stimulation periods (repeated measures ANOVA p<0.0001 with Tukey’s posthoc comparisons shown). **D)** Mean preferred firing phase (mu) of ChR2-tagged PV+ neurons in DG and CA1 of healthy PV-Cre mice during baseline (blue), trough-targeted stimulation (purple), and peak-targeted stimulation (pink; N=5 mice, n=25 cells). Peak-targeted stimulation significantly altered the distribution of preferred firing phases compared to baseline and trough-targeted stimulation periods (Kuiper test p≤0.001, circular k-test p<0.002 for both comparisons), but did not align them to the peak of theta (mean mu with peak stimulation: 203.5°; Rao test for non-uniformity p=0.50). **E)** Peak-targeted stimulation increased PV+ cell spiking at the peak of theta without impacting spiking at the originally preferred phase (the trough). **F)** The strength of phase preference, or r-value, was diminished during trough-targeted stimulation and peak-targeted stimulation periods (repeated measures ANOVA p=0.003 with Tukey’s posthoc comparisons shown).

**Fig. S6.**
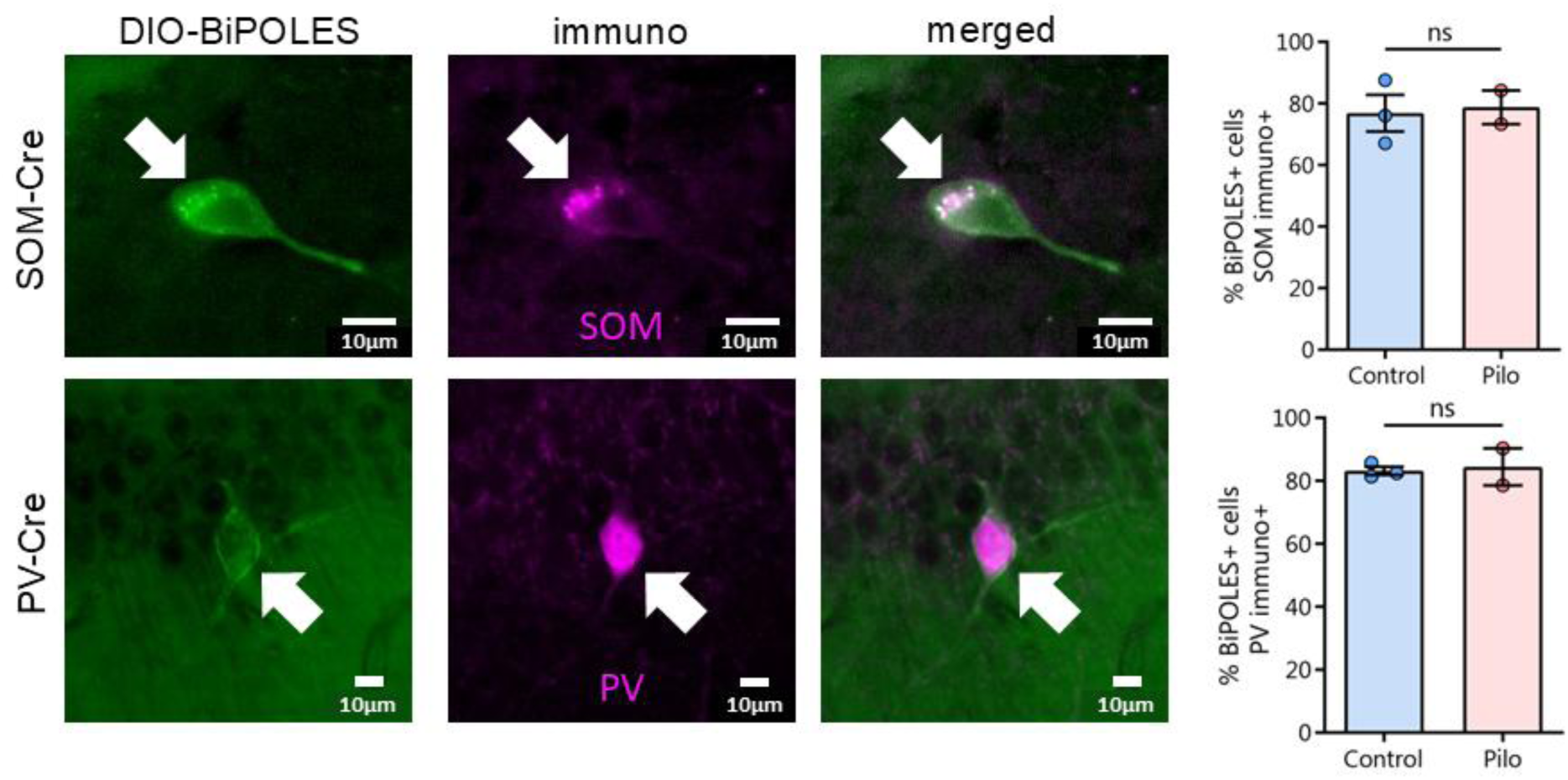
Validation of viral-transgenic approach to labeling SOM+ and PV+ cells in the dorsal hippocampus of control and pilocarpine-treated mice. Top: Cre-dependent somBiPOLES (AAV1-hSyn-DIO-somBiPOLES-mCerulean) expression colocalized with somatostatin (SOM) immunofluorescence in the dorsal hippocampus of control and epileptic SOM-Cre mice (unpaired t-test p=0.839). Bottom: Cre-dependent somBiPOLES expression colocalized with parvalbumin (PV) immunofluorescence in the dorsal hippocampus of control and epileptic PV-Cre mice (unpaired t-test p=0.806) Note that cell-type specific expression patterns were confirmed and were consistent across viruses (i.e., SOM-Cre mice showed expression of Cre-dependent viruses primarily in the hilus whereas PV-Cre mice showed expression of Cre-dependent viruses primarily at the granule cell layer-hilar border). Further, no Cre-dependent viral expression was seen in either control or epileptic C57BL6 mice [Viruses tested: AAV1-hSyn-DIO-somBiPOLES-mCerulean (N=3 control N=3 pilo), AAV1-hSyn-DIO-eGFP (N=3 control, N=3 pilo), AAV1-EF1a-DIO-hChR2(H134R)-eYFP-WPRE (N=4 control, N=6 pilo), AAV1-EF1a-DIO-eYFP (N=3 control, N=4 pilo)].

**Fig. S7:**
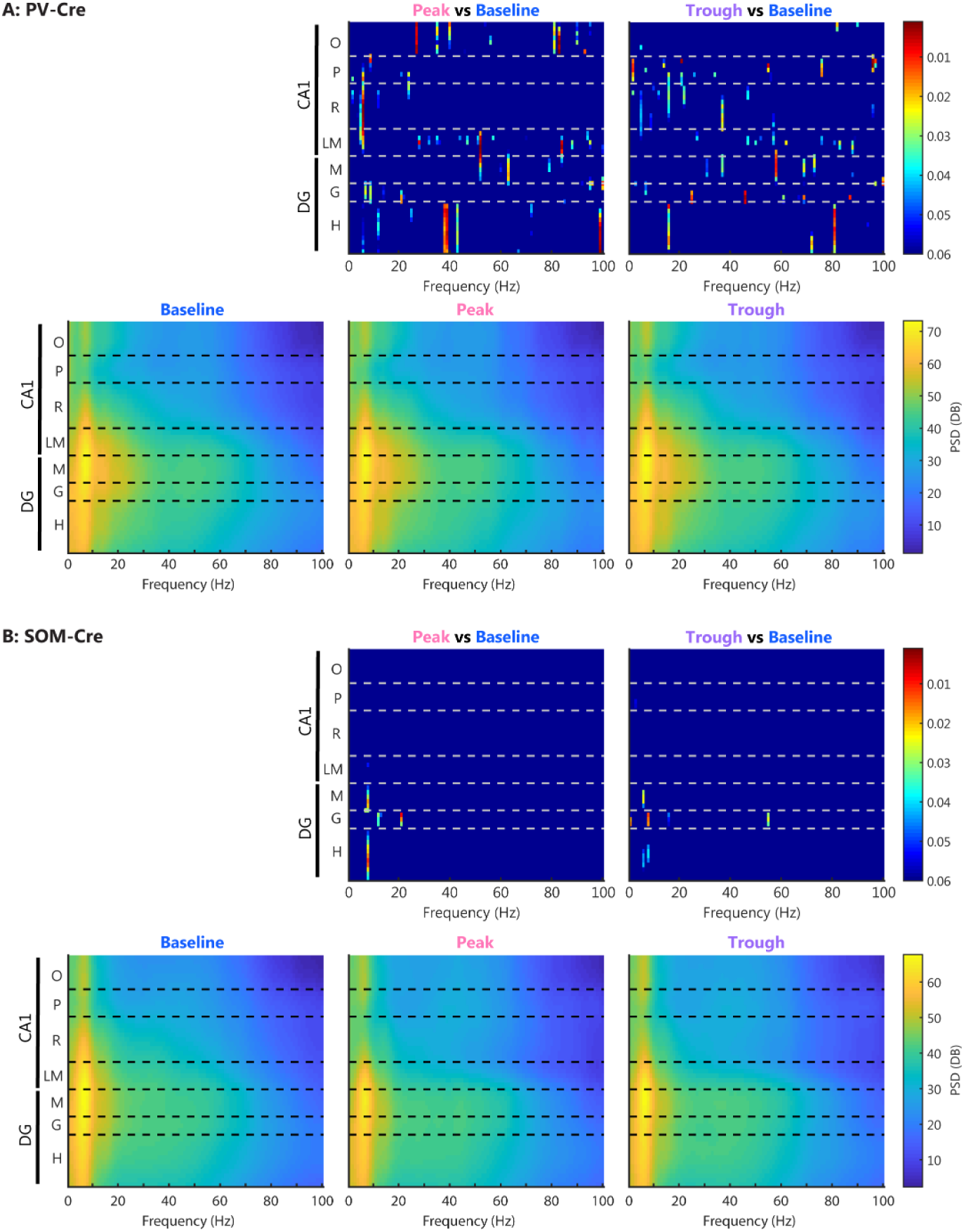
Manipulating DG PV+ or SOM+ cell theta phase locking does not alter oscillatory power in the dorsal hippocampus. **A-B** Power spectral density (PSD) plots from control PV-Cre (A) or SOM-Cre (B) mice injected with Cre-dependent BiPOLES in the DG. PSD plots show oscillatory power from 0-100Hz (1Hz bins) at 51 depth locations across CA1 and the DG. PSDs were compared using a t-test for each frequency-location comparison (5100 comparisons per stimulation phase per cell type) between running periods during baseline and peak-targeted stimulation or trough-targeted stimulation (with inhibitory light pulses delivered at the opposite phases). Heatmaps showing p-values from these t-tests are shown above the relevant PSD plot. Using a highly exploratory uncorrected alpha of 0.05, fewer comparisons than the expected 5% false positive rate emerged (with PV+ peak-stimulation, 3.94% of comparisons were significant; PV+ trough-stim 2.92%; SOM+ peak-stim 0.49%; SOM+ trough-stim: 0.39%, 10 PV-Cre mice, N=7 SOM-Cre). As a positive control, we confirmed that this method could easily detect changes in oscillatory power, such as those known to occur during locomotion (e.g., in PV-Cre mice baseline power during locomotion was normalized to power during immobility and compared in the same manner as above; these comparisons greatly exceeded the expected 5% false positive rate, with 77.16% of comparisons producing an alpha less than 0.05, N=14 mice).

**Fig. S8.**
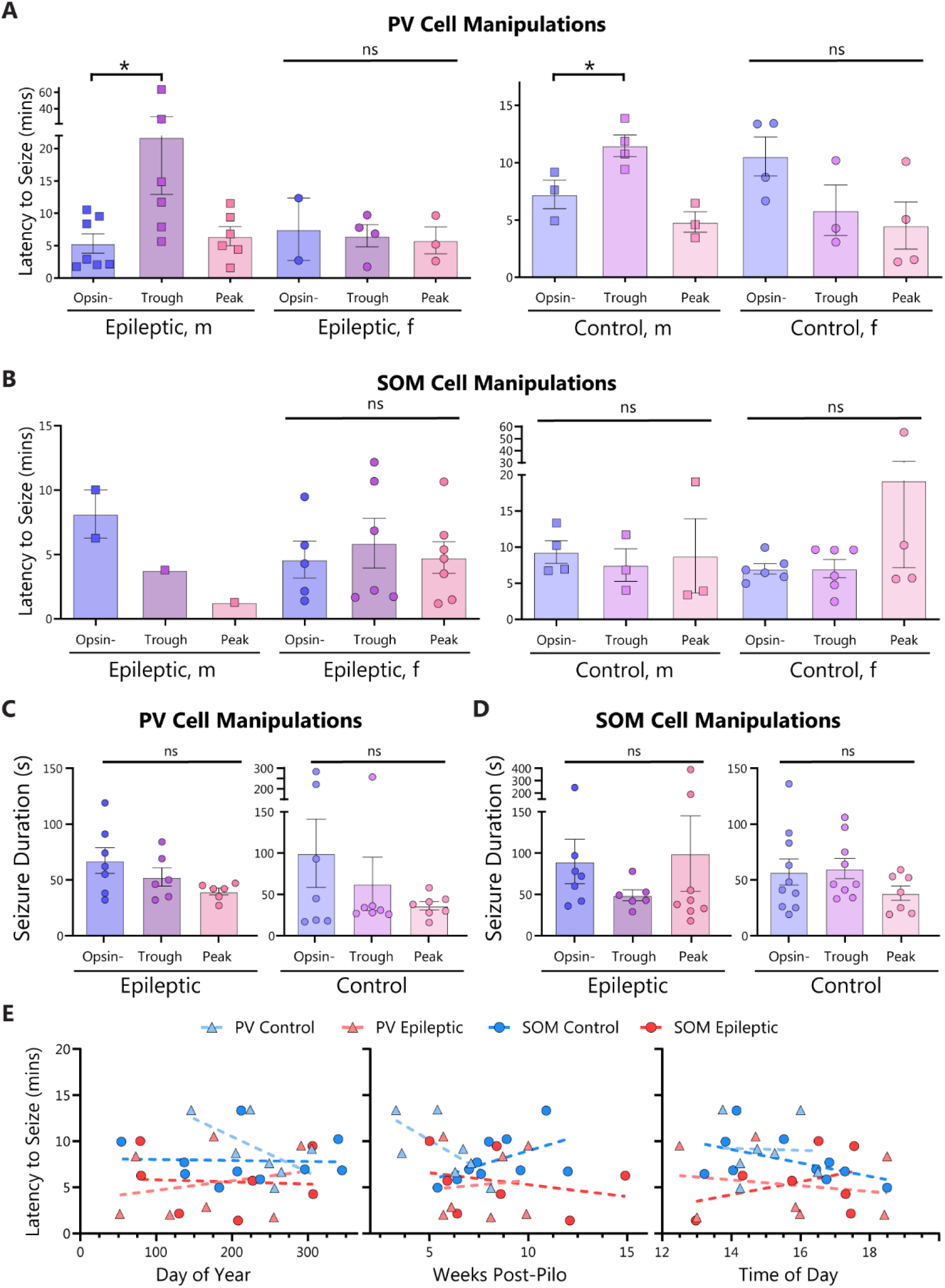
Exploring sources of variability in latency to seize and seizure duration following acute kainic acid injection. **A)** Latency to KA-induced seizure in epileptic (left) and control (right) PV-Cre mice, broken down by sex. Stimulating PV+ cells at the trough of CA1 theta increased time to seizure in both male control (one-way ANOVA p=0.006 with Dunnett’s multiple comparison test p=0.03) and male epileptic (Kruskal-Wallis ANOVA p=0.040 with Dunn’s multiple comparisons test p=0.042) mice, but not in female control (one-way ANOVA p=0.122) or female epileptic (one-way ANOVA p=0.909) mice. Note that the Epileptic male data is reproduced in Figure 4. **B)** Latency to KA-induced seizure in epileptic (left) and control (right) SOM-Cre mice, broken down by sex. There were no significant differences in latency to seize (one-way ANOVAs, all p>0.05), but note the low sample sizes of male epileptic SOM-Cre mice. **C)** KA-induced seizure duration was not altered when PV+ cell phase locking was forced to the trough or peak of CA1 theta in epileptic (left, males only, one-way ANOVA p=0.112) or control (right, combined sexes, Kruskal-Wallis ANOVA p=0.892) mice. Note there was also no change in seizure duration in the epileptic female mice (not shown, one-way ANOVA p=0.611). **D)** KA-induced seizure duration was not altered when SOM cell phase locking was forced to the trough or peak of CA1 theta in epileptic (left, one-way ANOVA p=0.575) or control (right, one-way ANOVA p=0.302) mice (sexes combined). **E)** No correlations between the latency to seizure in opsin-negative animals and day of the year, weeks post-pilo, or time of day (simple linear regressions, all p>0.05). Note that only male animals are included in the PV epileptic group, to match what is shown in Figure 4, but none of the results were changed by including the N=2 female PV epileptic opsin-animals.

**Table S1.** Summary of p-values comparing DG inhibitory theta phase locking in male and female control and epileptic mice. These statistics accompany Figure S2A. Note that the Watson-Williams and circular k-tests were performed post-hoc only if the Kuiper test showed a significant difference. * p<0.05, ** p≤0.01, *** p≤0.001, ****p≤0.0001.

|  | Kuiper test | Watson-Williams test | Circular k-test |
| --- | --- | --- | --- |
| Female Control vs Female Pilo | *** $p \leq 0.001$ | *** $p = 0.0005$ | *** $p = 0.0002$ |
| Male Control vs Male Pilo | *** $p \leq 0.001$ | ** $p = 0.0013$ | **** $p < 0.0001$ |
| Female Control vs Male Control | NS: $p > 0.1$ | N/A | N/A |
| Female Pilo vs Male Pilo | ** $p \leq 0.010$ | **** $p < 0.0001$ | NS: $p = 0.158$ |

## ACKNOWLEDGEMENTS

We would like to wholeheartedly thank the Shuman and Cai lab groups for providing their support and thoughtful feedback throughout the course of this project. In particular, we would like to thank Alia Abdelhameed and Helen Liu for assistance with animal training and tissue processing, Lucia Page-Harley for methodological set-up and support, and Lingxuan Chen, Nadia Khan, William Mau, Zachary Pennington, Justin Lines, Yosif Zaki, and Brian Sweis for helpful conversations that shaped this project. Corin Humphrey passed away in October 2020 after making substantive methodological contributions to this project. She agreed to be an author on this project before her death.

## Author contributions

Conceptualization: ZCW, PAP, TS

Methodology: ZCW, PAP, YF, LMV

Investigation: ZCW, CK, SIL, EK, KEG, GCD, CH

Formal Analysis: ZCW, PAP, CK

Software: PAP

Visualization: ZCW, PAP, TS

Funding acquisition: ZCW, PAP, YF, LMV, DJC, TS

Project administration: ZCW, TS

Supervision: ZCW, TS

Writing – original draft: ZCW, PAP, CK

Writing – review & editing: ZCW, PAP, CDA, DJC, TS

## Data and materials availability

All software and hardware requirements to implement real-time phase manipulations during behavior are available online (https://github.com/ShumanLab/PhaSER). All LFP and spike data will be made publicly available upon publication. Any additional information required to re-analyze the data reported in this paper is available from the lead contact upon request.

